# Evolution of a Plasmid Regulatory Circuit Ameliorates Plasmid Fitness Cost

**DOI:** 10.1101/2024.02.05.579024

**Authors:** Clinton A. Elg, Erin Mack, Michael Rolfsmeier, Thomas C. McLean, Olivia Kosterlitz, Elizabeth Soderling, Solana Narum, Paul A. Rowley, Christopher M. Thomas, Eva M. Top

## Abstract

Plasmids play a major role in rapid adaptation of bacteria by facilitating horizontal transfer of diverse genes, most notably those conferring antibiotic resistance. While most plasmids that replicate in a broad range of bacteria also persist well in diverse hosts, there are exceptions that are poorly understood. We investigated why a broad-host range plasmid, pBP136, originally found in clinical *Bordetella pertussis* isolates, quickly became extinct in laboratory *Escherichia coli* populations. Through experimental evolution we found that inactivation of a previously uncharacterized plasmid gene, *upf31*, drastically improved plasmid maintenance in *E. coli*. This gene inactivation resulted in decreased transcription of the global plasmid regulators (*korA*, *korB,* and *korC)* and numerous genes in their regulons. It also caused transcriptional changes in many chromosomal genes primarily related to metabolism. *In silico* analyses suggested that the change in plasmid transcriptome may be initiated by Upf31 interacting with the plasmid regulator KorB. Expression of *upf31 in trans* negatively affected persistence of pBP136Δ*upf31* as well as the closely related archetypal IncP-1β plasmid R751, which is stable in *E. coli* and natively encodes a truncated *upf31* allele. Our results demonstrate that while the *upf31* allele in pBP136 might advantageously modulate gene expression in its original host, *B. pertussis*, it has harmful effects in *E. coli*. Thus, evolution of a single plasmid gene can change the range of hosts in which that plasmid persists, due to effects on the regulation of plasmid gene transcription.

## INTRODUCTION

Bacterial evolution and rapid adaptation are frequently shaped by the horizontal transfer of genes, including those transferred by extra-chromosomal DNA elements known as plasmids (Norman et al. 2009; Soucy et al. 2015). One example is the plasmid-encoded spread of antibiotic resistance, which enables bacterial pathogens to resist traditional therapeutic antibiotic treatments, leading to the rise of highly resistant “superbugs” that increasingly threaten human health (Murray et al., 2022; San Millan, 2018). Understanding how plasmids evolve to successfully transfer and maintain themselves in bacterial populations and communities is thus a critical step towards limiting the spread of antibiotic resistance.

A central metric for assessing the degree to which a plasmid remains in a bacterial population over time is termed ‘plasmid persistence’. Plasmid-host pairs often differ in plasmid persistence, and even the same plasmid has been shown to persist differently in closely related species (De Gelder et al. 2007; Kottara et al. 2018). Instances of poor plasmid persistence can often be attributed to decreased fitness of the plasmid-containing bacterium relative to its plasmid-free counterpart. This fitness cost of plasmid carriage imposed on the bacterial host is due to several factors, including metabolic costs and molecular conflicts between plasmid and host machinery (Modi and Adams, 1991). Several studies have shown compensatory mutations that arise during serial batch cultivation of plasmid-host pairs; these mutations ameliorate plasmid fitness cost and thereby restore plasmid persistence. Some of the important early studies showed that such cost amelioration could occur through genetic changes on the plasmid but were unable to explore the underlying molecular mechanisms (Bouma and Lenski, 1988; Dahlberg and Chao, 2003). With the advent of next generation sequencing, compensatory mutations have been identified in specific plasmid genes, including those encoding accessory functions (Bottery et al. 2017; Stalder et al. 2017), replication initiation (Sota et al. 2010; Hughes et al. 2012; Yano et al. 2016), and conjugation machinery (De Gelder et al. 2008; Jordt et al. 2020; Porse et al. 2016; Yang et al. 2023). Additionally, chromosomal evolution can stabilize host-plasmid pairs through amelioration of plasmid cost via mutations targeting global regulators (Harrison et al. 2015; Stalder et al. 2017) and putative helicases (San Millan et al. 2014; Loftie-Eaton et al. 2017). Thus, compensatory mutations that increase plasmid persistence can occur on the plasmid, chromosome, or both (Hall et al. 2021).

Some plasmid types, such as those of the incompatibility group IncP-1 used in this study, are known to be highly persistent across many Proteobacteria (Schmidhauser and Helinski, 1985: Shintani et al. 2010; Yano et al. 2012, 2013; Jain and Srivastava, 2013; Klümper et al. 2015;), with few exceptions (De Gelder et al. 2007; Kottara et al. 2018). This remarkable persistence is aided by several conserved regions (Thorsted et al. 1998; Thomas, 2000; Sen et al. 2013) of backbone genes, which ensure the fidelity of critical functions like replication, stable inheritance and control, mating pair formation and conjugative DNA transfer. These backbone genes are controlled by a complex genetic regulatory circuit with four repressors located chiefly within the stable inheritance and control region (Bingle et al. 2005; Rajasekar et al. 2016). The regulators are part of a negative feedback loop referred to as autogenous control (Bingle and Thomas, 2001) that limits their own expression and that of the operons they control. This in turn allows fine-tuned and temporal plasmid gene expression to minimize cost of plasmid carriage to the host and prevent potential conflicts with host cellular machinery. Collectively, this regulatory network enables IncP-1 plasmids to replicate, transfer, and be stably maintained in multiple classes of the Proteobacteria (Schmidhauser and Helinski, 1985; Shintani et al. 2010; Yano et al. 2012, 2013); Jain and Srivastava, 2013; Klümper et al. 2015).

Given the known broad host range of IncP-1 plasmids and their role in the spread of antibiotic resistance (Popowska and Krawczyk-Balska, 2013) we sought to examine the previously reported but unexplained poor persistence of subgroup IncP-1β plasmid pBP136Km in several *Escherichia coli* strains (Sota and Top, 2008). This plasmid was previously obtained by inserting a kanamycin resistance gene in pBP136 (Sota et al. 2007), which was found in a *Bordetella pertussis* strain isolated from a lethal infection in an infant in Japan (Kamachi et al. 2006).

We report that a single genetic change in pBP136Km that occurred within 50 generations of evolution in *E. coli* in the absence of antibiotic selection drastically increased the plasmid’s persistence. This improved persistence was explained by a decrease in the plasmid fitness cost due to partial deletion and presumed inactivation of a plasmid-encoded gene with unknown protein function, *upf31*. This gene was previously thought to encode a putative adenine methylase that is associated with an intriguing set of clustered repeats specific to the IncP-1β plasmids (Thorsted et al. 1998). The high cost of plasmid carriage caused by this *upf31* allele was shown to extend to the archetypal IncP-1β plasmid R751 (Thorsted et al. 1998), which is otherwise stable in *E. coli*. In pBP136Km the presence of Upf31 was associated with changes in expression of plasmid backbone regulatory genes and chromosomal genes, which we propose may be due to its physical interaction with plasmid regulator KorB. Our study highlights that evolution of the intricate gene regulatory systems of self-transmissible plasmids can improve plasmid-bacteria pairings by amelioration of plasmid fitness costs.

## RESULTS

### Plasmid-encoded *upf31* results in poor plasmid persistence in *E. coli* hosts

We first sought to confirm a previous finding (Sota and Top, 2008) that plasmid pBP136Km shows poor persistence in various *E. coli* hosts. Using *E. coli* K-12 MG1655 (hereafter K-12), we performed a plasmid persistence assay entailing daily passage of triplicate cultures in non-selective conditions (*i.e.* without antibiotics). This initial assay with the ancestral host-plasmid pair showed poor plasmid persistence (Figure 1a in blue). In line with the previous study, we randomly isolated one plasmid-bearing clone from each of the triplicate populations on Day 5 and tested plasmid persistence in these clones. The persistence of the plasmid in these clones was markedly improved (clones A, B, and C in Figure 1a in red, green, magenta, respectively).

**Figure 1:**
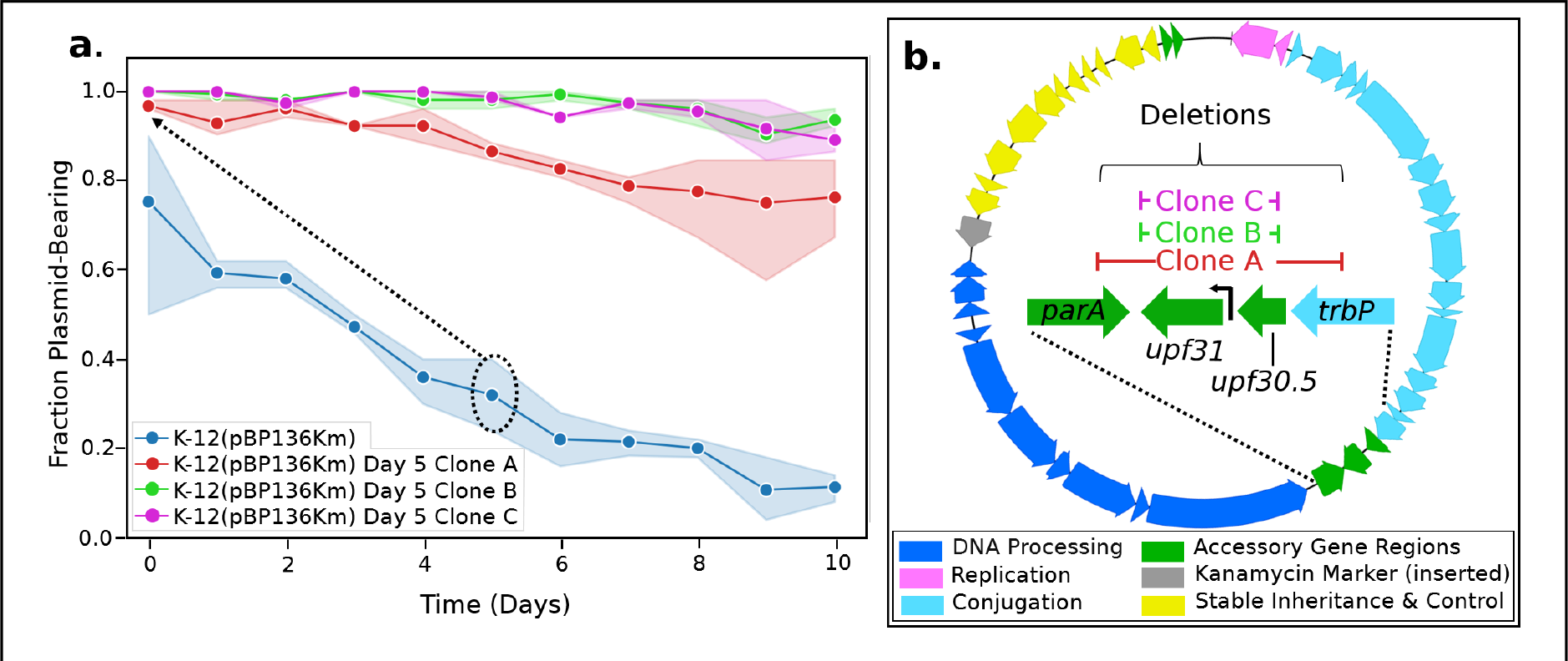
Plasmid evolution rapidly improved pBP136Km persistence. **(a)** Persistence of the ancestral plasmid pBP136Km (blue) and three populations (red, green, magenta) inoculated with clones that were isolated on Day 5 of the ancestral plasmid assay (see dotted ellipse and arrow). Clones isolated on Day 5 had greatly improved persistence compared to the ancestor. Lighter shades indicate 95% confidence interval. **(b)** Genomic map of pBP136Km in evolved Day 5 clones revealed deletions in the accessory region (green) between the *trb*-*tra* operons (light and dark blue, respectively).

To identify the genetic changes that may explain the observed change in plasmid persistence, we completely sequenced the three isolated clones (Supplemental Table S1). We found deletions within pBP136Km consistently targeting the region often associated with accessory genes between the *tra* and *trb* operons, and clones B and C had identical plasmids (Figure 1b). Additionally, no genetic changes were observed in the chromosomes of these three clones. As the plasmid deletions in all three clones at least included *upf30.5* and *upf31.0*, we suspected the presence of one or both genes to be responsible for the poor plasmid persistence in *E. coli*.

We also routinely sequenced clones recovered after transferring the ancestral plasmid between strains of *E. coli* by conjugation (*i.e.* matings) or electroporation. Among these clones were several with deletions either internal to *upf31* or in its putative promoter (Supplemental Table S1). To test if deletion of *upf31* alone could explain improved plasmid persistence, we compared the persistence of one of these recovered variants, named pBP136KmΔ*upf31,* to that of the ancestral plasmid pBP136Km in the same host, *E. coli* K-12 (Figure 2a).

**Figure 2:**
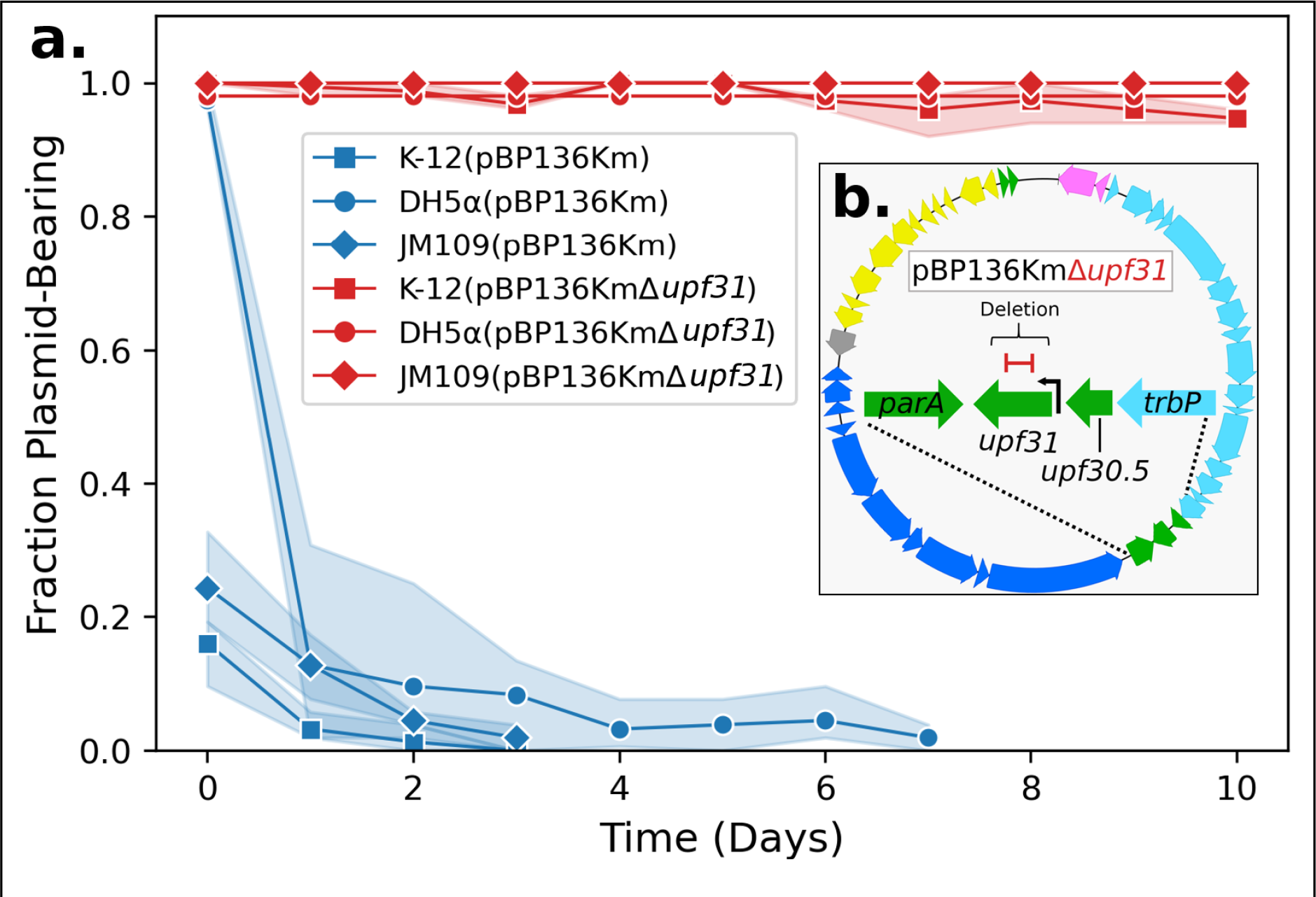
*upf31* inactivation alone explained improved persistence in *E. coli* hosts. **(a)** The poor persistence of plasmid pBP136Km (blue lines) in *E. coli* strains K-12 MG1655, DH5α, and JM109 (square, circle, and diamond respectively) was markedly improved after an internal 168-bp deletion in plasmid gene *upf31* (red lines). Lighter shades indicate 95% confidence interval. **(b)** A map of pBP136KmΔ*upf31* showing the 168-bp deletion (red) within the coding sequence of *upf31*.

This selected plasmid contained a 168-base pair (bp) deletion internal to the open reading frame of *upf31* whereas *upf30.5* was intact (Figure 1b and Supplemental Table S1). Given that the plasmid with inactivated *upf31* was persistent for at least 10 days, this gene of unknown function must be responsible for the very poor persistence of pBP136 (compare red versus blue squares of Figure 2a). The improvement in plasmid persistence after inactivation of *upf31* was also seen in other well-studied strains of *E. coli* including DH5α and JM109 (Figure 2a).

Importantly, the pBP136KmΔ*upf31* containing clone was isolated from an overnight mating followed by plating for transconjugants on selective media (Supplemental Table S1). This provides an example of very rapid plasmid evolution that results in remarkably improved plasmid persistence (*i.e.* < 48 hours, see Hall et al. 2020).

### Host fitness was lowered in the presence of upf31 and pBP136Km

To investigate if the improved plasmid persistence associated with *upf31* inactivation was due to amelioration of plasmid fitness costs, we employed two approaches: *1)* a comparison of the growth dynamics between the two plasmid-host pairs, with a focus on the maximum growth rate and carrying capacity, and *2)* competition assays involving plasmid-bearing and plasmid-free strains. First, we observed that K-12 containing pBP136KmΔ*upf31* had a significantly higher maximum growth rate and reached a higher final density in comparison to K-12 containing ancestral pBP136Km (Figure 3a). These results demonstrate that the presence of *upf31* encoded on pBP136Km imposes a substantial fitness cost on its host.

**Figure 3:**
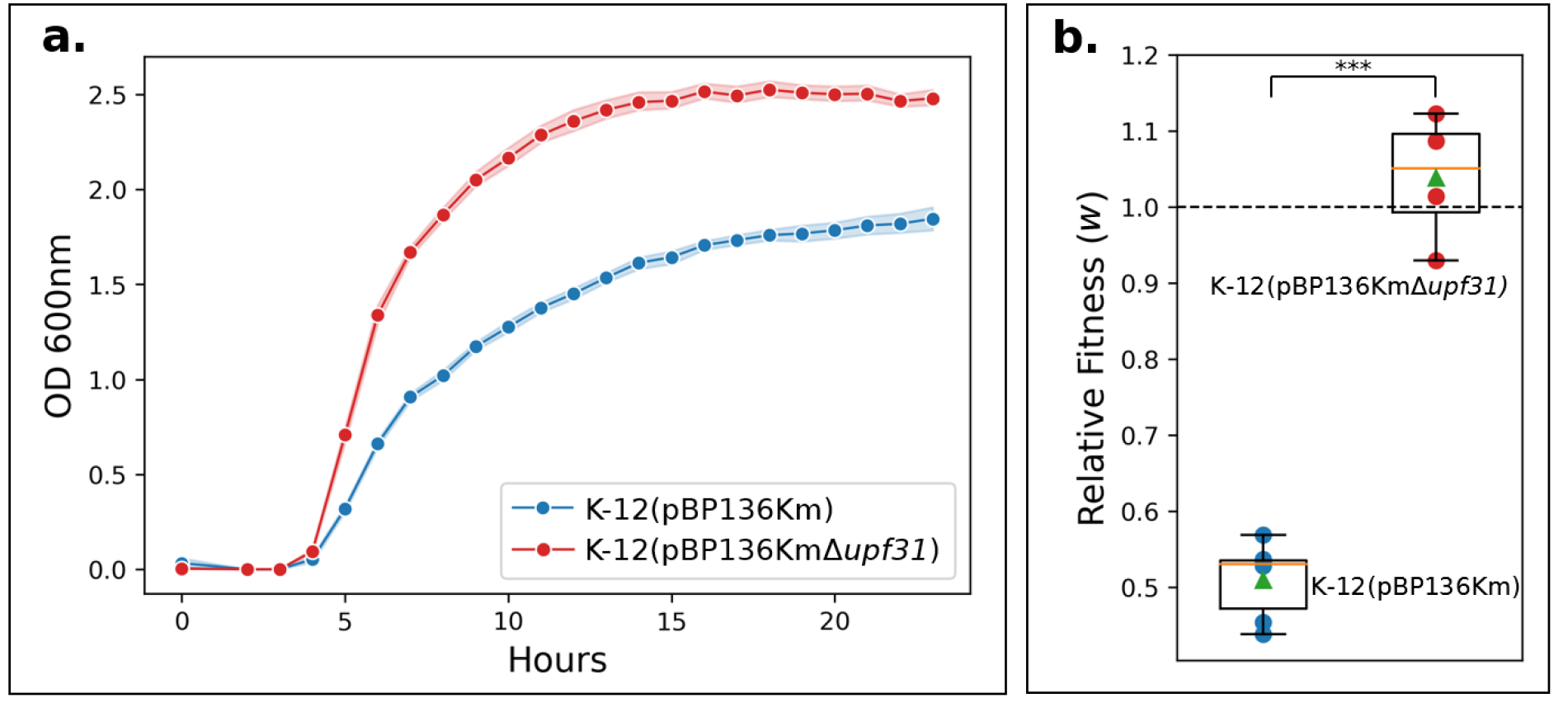
The presence of *upf31* encoded on pBP136Km lowered host fitness. **(a)** Bacterial growth in batch culture was slower (blue) when carrying pBP136Km with ancestral *upf31* compared to the evolved plasmid pBP136KmΔ*upf31* with inactivated *upf31* (red). Lighter shades indicate 95% confidence interval. (Two sample independent T-test of max growth rates, *n*=10, p-value= 2.35x10-3) **(b)** With the fitness of plasmid-free K-12 normalized to 1 (dashed line), the relative fitness (*w*) of *E. coli* with ancestral pBP136Km (in blue) was 0.51, *i.e.,* a ∼49% cost (Two sample independent T-test, *n*=5, p-value=1.70x10-6). The relative fitness of evolved genotype K-12 (pBP136KmΔ*upf31*) to plasmid-free K-12 was statistically indistinguishable (Two sample independent T-test, *n*=5, p-value=0.40). Box is interquartile range, green triangle is mean, orange line is median, ***=p ≤ 0.001.

Next, we confirmed the results of the growth rate observations by estimating the cost of the ancestral and evolved plasmid in competition assays. This was done by individually competing each plasmid-bearing strain against the plasmid-free K-12. We note that direct competition assays between identical hosts with and without a highly transmissible conjugative plasmid such as pBP136Km can be confounded by plasmid transfer during the assays. While IncP-1 plasmids are less efficient at transferring in liquid than on surfaces, they have been shown to transfer at detectable rates (Zhong et al. 2010). To overcome this limitation, we developed a novel low-density competition assay which allows competing two strains at densities too low for appreciable plasmid transfer to occur (see Methods). Indeed, as conjugation requires cell contact, decreasing initial densities results in decreasing cell collisions and thus plasmid transfer events (Kosterlitz et al. 2022). The results (Fig 3b) show a fitness of K-12 (pBP136Km) relative to K-12 of only 0.51, suggesting the plasmid caused a 49% reduction in fitness. By contrast, the fitness of K-12 (pBP136KmΔ*upf31*) was statistically indistinguishable from that of K-12. We conclude that the persistence differences in Figure 2 can be attributed to a very large fitness cost conferred by pBP136Km, which was alleviated by the inactivation of one plasmid gene, *upf31*.

### The fitness cost imposed by *upf31* requires the presence of pBP136Km

We next sought to understand if the expression of *upf31* directly imposed a fitness cost on K-12 in the absence of plasmid pBP136Km. This was done by expressing *upf31* within K-12 *in trans* from expression vector pCW-LIC-*upf31.* Production of protein Upf31 was verified via induction of pCW-LIC-*upf31* and observation of the associated 25.4 kDa product (Supplemental Figure S1). As a negative control we used the non-modified vector pCW-LIC-*sacB*. The maximum growth rates of K-12 with or without the Upf31 protein were not significantly different (Fig 4a). This suggests that Upf31 does not impose a significant fitness cost on K-12 in the absence of pBP136Km where it is naturally encoded. In contrast, when *upf31* was expressed *in trans* in the presence of pBP136KmΔ*upf31,* the maximum growth rate was significantly reduced compared to the same strain with the control vector (Fig 4b). Thus, we demonstrated that Upf31 did not directly impose a fitness cost on K-12 but rather the interaction of Upf31 with pBP136Km was responsible for the host fitness cost.

**Figure 4:**
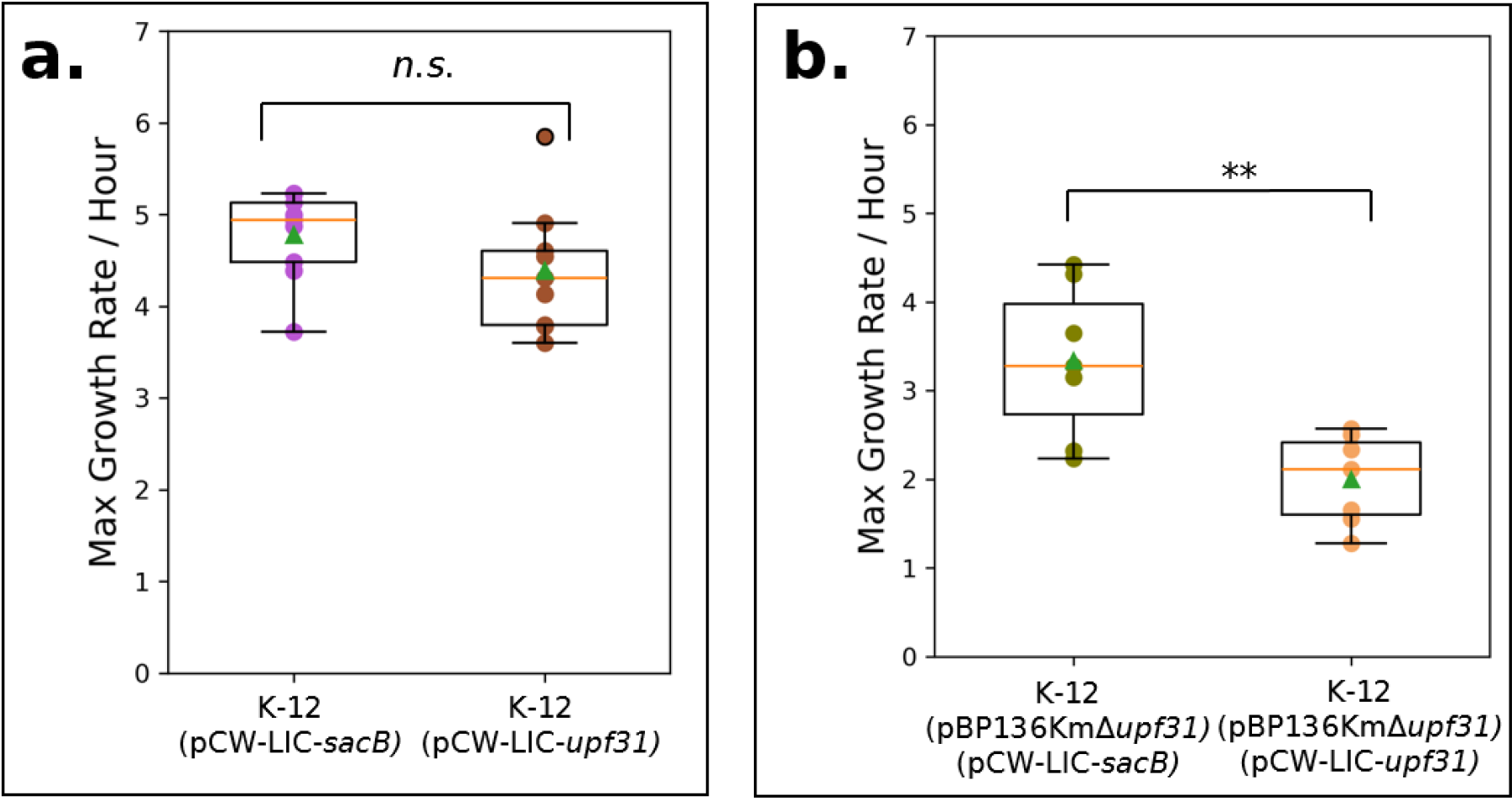
Upf31 requires the presence of pBP136Km to significantly reduce K-12 growth rate. **(a)** In the absence of plasmid pBP136Km there was no significant difference in maximum growth rate between *E. coli* K-12 expressing control *sacB* (purple) and *upf31* (brown) (Two sample independent T-test, *n*=10, p-value=0.46). **(b)** K-12 (pBP136KmΔ*upf31*) with *upf31* expressed *in trans* (orange) showed a lower maximum growth rate than K-12 (pBP136KmΔ*upf31*) with control *sacB* expressed *in trans* (green) (Two sample independent T-test*, n*=10, p-value=4.23x10-3). Box is interquartile range, green triangle is mean, orange line is median, *n.s.* = p > 0.05, **=p ≤ 0.01.

### Gene *upf31* also effects persistence and fitness cost of archetype Inc-1β plasmid R751 when complemented *in trans*

We examined if the high cost of plasmid carriage observed with *upf31* extends to other IncP-1β plasmids. The archetype IncP-1β plasmid R751(Thorsted et al. 1998) encodes a *upf31* homolog with a 66 base pair deletion at the 3’ end compared to pBP136Km. The high persistence of R751 in *E. coli* suggests that this allele does not negatively affect *E. coli* fitness (Supplemental Figure S2). To test if *upf31* from pBP136Km affects K-12 (R751) when expressed *in trans*, we compared K-12 (R751) (pCW-LIC-*upf31*) to control K-12 (R751) (pCW- LIC-*sacB*) in terms of plasmid persistence and growth rate. Plasmid R751 showed lower persistence and resulted in a statistically significant lower host growth rate (Supplemental Figure S2) when pBP136Km’s *upf31* is present. This suggests that the longer *upf31* allele of pBP136 has a negative effect on R751 cost and persistence.

Forty-nine IncP-1β plasmids were identified in NCBI that contain a gene with full-length homology to the ancestral *upf31* and associated promoter in pBP136, although none have an identical protein sequence (Supplemental Data S1). We tested the persistence of two of these plasmids, pAKD1, isolated from forest soil (Drønen et al. 1998), and the wastewater associated pALTS29 (Law et al. 2021). Whereas plasmid pAKD1 showed poor persistence, similar to pBP136Km, pALTS29 demonstrated high persistence (Supplemental Figure S3). The *upf31* gene product of plasmid pAKD1 differs from that of pBP136Km in ten amino acids, with three of these being non-conservative substitutions (D76A, A91V, R124G – see Supplemental Table S2). Upf31 from pALTS29 differed from that of pBP136Km in twelve amino acids, including the same three non-conservative substitutions found in pAKD1. When comparing the Upf31 amino acid sequence between all three plasmids, pAKD1 had a unique R60Q change and pALTS29 had three unique differences in L78R, Q82P, S86T. Plasmid pALTS29 may be worth further analysis, especially since the wastewater environment in which it was found might have selected plasmid variants that persist better in a range of *Enterobacteriaceae*.

### The computationally predicted methylase function of Upf31 was not supported by experimental results

To provide further insight into the possible function(s) of Upf31 we performed a bioinformatic analysis using Phyre2. The output strongly predicted Upf31 to be a Dam methylase homolog with 70% of the amino acids modeling with 100% confidence (Supplemental Data S2). This is consistent with the pBP136 reference genome including a computationally automated annotation of *upf31* as encoding a DNA methylase based on protein similarity as noted in the original annotation of archetypal IncP-1β plasmid R751 (Thorsted et al. 1998). To test if Upf31 had Dam activity, we digested genomic DNA extracted from *dam^-^* strain *E. coli* JM110 with and without pBP136Km and *dam^+^* K-12, using restriction enzymes that digest only methylated or non-methylated ‘GATC’ DNA sequences. The results suggest that Upf31 is not a functional Dam homolog as the restriction profiles of DNA from JM110 (pBP136Km) were opposite those of the *dam+* strain K-12 (Supplemental Figure S4).

Next, to determine if Upf31 more broadly encoded a methyltransferase, we used the base pair modification detection available via Single Molecule Real Time (SMRT) sequencing. This method has the benefit of not requiring *a priori* knowledge of the base pair modification chemistry or target DNA sequence of the putative methylase. Briefly, the methylation-free *E. coli* ER2796 (Anton et al. 2015) was used with *upf31* expressed from vector pCW-LIC-*upf31* or pCW-LIC-*sacB* as the vector control. Genomic DNA extracts of both strains were analyzed using the base modification analysis on a Pacbio Sequel II, which measures the kinetic incorporation of base pairs to detect patterns distinctive from non-methylated base pairs. The output showed no discernible kinetic patterns associated with methylated base pair incorporation, indicating that Upf31 does not function as a methylase under our experimental conditions, in contrast to the computational predictions.

### The presence of *upf31* alters the expression of global plasmid regulators and associated operons

To understand how *upf31* and pBP136Km interact to influence *E. coli* fitness, we next performed RNA-seq on K-12 containing pBP136Km and K-12 containing pBP136KmΔ*upf31*. We considered K-12 (pBP136Km) to be the treatment (presence of the intact gene *upf31*), and K- 12 (pBP136KmΔ*upf31*) with the internal 168-bp deletion to be the control (absence of a functional *upf31*). The presence of *upf31* led to greater than two-fold expression changes in 654 of the ∼4,400 chromosomal genes, with 412 of those upregulated and 242 downregulated (adjusted P-value < 0.05, Supplemental Data S3). The gene ontology tool DAVID (Dennis et al. 2003) predicted downregulation of cellular pathways related to nitrogen and sugar metabolisms, while biosynthesis of siderophores, ABC transporters, and sulfur metabolism pathways were all upregulated (Supplemental Data S4).

Most interestingly, K-12 (pBP136Km) showed greater than two-fold expression increase in 11 of its 46 plasmid genes relative to the *upf31* deletion variant (adjusted P-value < 0.05, Figure 5a and Supplemental Data S3). These genes were primarily part of the stable inheritance and control region made up of regulons controlled by proteins KorA, KorB and KorC (Figure 5a in yellow). Strikingly, K-12 (pBP136Km) showed a relative decrease in expression in only one gene, *upf31* itself. This relative comparison of gene expression was made possible by pBP136KmΔ*upf31* not having a complete *upf31* deletion but rather an internal 168-bp deletion that left an inactivated 507 bp *upf31* product to be expressed (Figure 5b).

**Figure 5:**
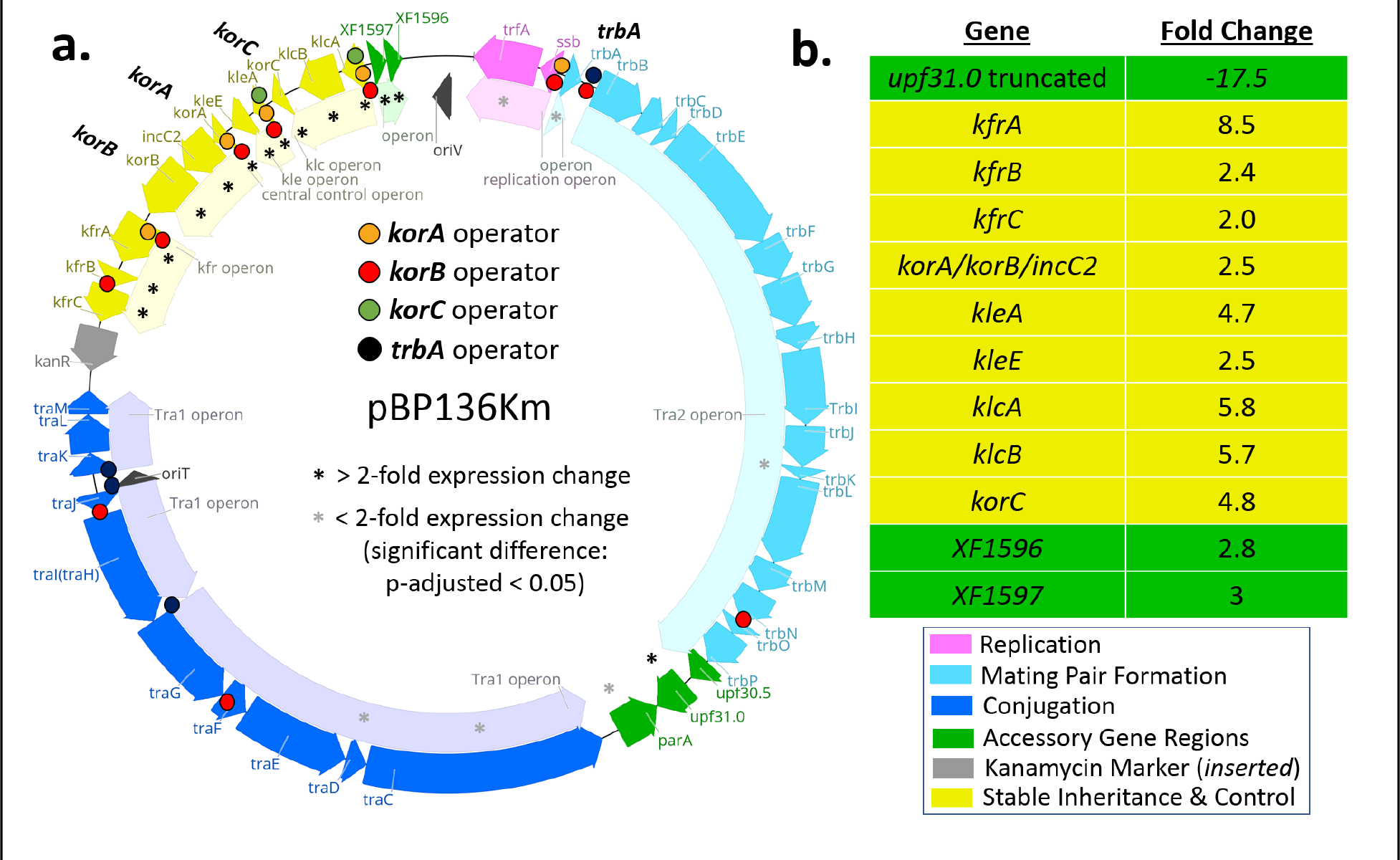
Expression of global plasmid regulators and their associated operons was increased in the presence of ancestral *upf31* compared to the deletion mutant. **(a)** Genetic map of pBP136Km with operons and operator sites annotated for global regulators KorA, KorB, KorC, and TrbA. Asterisks denote differentially expressed plasmid genes in the presence of *upf31*. **(b)** Table of plasmid-encoded genes with at least 2-fold differential expression (black asterisks in panel a) in the presence of *upf31.* Note expression of the ancestral *upf31* is ∼16-fold lower than that of the truncated *upf31* knockout.

The effect of Upf31 to cause at least two-fold increased expression of all four of the canonical IncP-1 regulatory genes (Adamczyk and Jagura-Burdzy, 2003)– *korA, korB, korC,* and *trbA* – along with the other genes in the operons they regulate (Figure 5a in bold) is notable.

These regulators are known to autogenously repress their own expression (Bingle and Thomas, 2001) by binding cognate operators near their respective promoters (orange, red, green, and black dots in Figure 5a). They are also known to repress promoters for the other operons in their regulons (yellow in Figure 5a). The increased repressor gene expression should therefore decrease rather than increase expression of all other regulated operons as the expression of the repressor genes increases. This suggests that *upf31* has a general effect of diminishing the repressive ability of *kor* regulatory genes, both in terms of their autogenous control and repression of their respective regulons.

### Upf31 represses its own transcription

A possible mechanistic explanation for the decrease in *upf31* transcription in the presence of Upf31 is that Upf31 represses its own transcription, like the previously IncP-1 plasmid regulators KorA, KorB, KorC and TrbA studied (Pansegrau et al. 1994). Testing this hypothesis requires first verifying the putative promoter upstream of upf31(*i.e.* upf31p) and then measuring expression from this promoter with and without Upf31 present in the cells. To test the functionality of the putative *upf31* promoter upf31p, a 68-bp region upstream of *upf31* was inserted into *xylE* reporter plasmid pGCMT1 to create pGCMTupf31p (Supplemental Figures S5 and S6). The *xylE* assay confirmed that this region possesses significant promoter activity with 7.4 XylE units compared to the 0.09 units for the empty vector (Supplemental Table 3).

Since the Thomas lab had previously failed to demonstrate strong repressor activity from the 3’-end truncated *upf31* in R751(Akhtar, 2002), we tested if adding the C-terminus of pBP136’s Upf31 to R751’s Upf31 restored the protein into a functional repressor. To do this, we constructed an R751/pBP136 hybrid *upf31* gene (aa 1 to 172 from R751 and aa 173 to 225 from pBP136) and inserted this hybrid gene into pBR322 to create pBR*upf31*. Expression from the *upf31* promoter on pGCMTupf31p was then measured in the presence and absence of pBR*upf31*. The full-length hybrid Upf31 expressed from pBR*upf31* caused repression of upf31p, resulting in an average of 0.09 XylE units compared to 4.76 XylE units with pBR322 as negative control (Supplemental Table 3). This confirmed that *upf31* expression is likely to be autoregulated via Upf31 binding near upf31p. The most obvious operator-like sequence in the region of this promoter consists of inverted 5’-CAGCATCG-3’ repeats, which belongs to the family of inverted repeats (IR) that occur in four groups across the IncP-1β genome (Thorsted et al. 1998). The presence of these strikingly conserved sequences of these IR across IncP-1β plasmids suggests that Upf31 may bind to similar sequences across IncP-1β plasmids. In conclusion, these assays confirmed the transcriptome results that suggested Upf31 represses itself and may have additional regulatory functions.

### Upf31 is computationally predicted to bind KorB via a C-terminal dimerization domain

A hypothesis to explain our transcriptome results emerges from the fact that the high level of repression associated with the stable inheritance and control region depends on the cooperative interaction of KorA (and TrbA) with KorB (Zatyka et al. 1997; Kostelidou et al. 1999; Bingle et al. 2008). If Upf31 were able to disrupt this cooperativity then it might decrease repression while allowing the concentration of the repressors to rise. To explore the hypothesis that Upf31 disrupts the cooperative interaction of KorA (and TrbA) with KorB we used Alphafold2 (Jumper et al. 2021) and ColabFold (Mirdita et al. 2022) to predict the structures of every individual pBP136 coding region in complex with Upf31. We subsequently ranked these interactions using a self-assessment ranking score (ranking_confidence: 0.2pTM + 0.8ipTM, see Supplemental Data S5). The structure prediction for Upf31 is that it possesses a globular DNA binding domain (residues 1 to 174) and a C-terminal dimerization domain (CTD) which is consistent with its proposed autogenous regulation of its own expression. However, the most exciting result of this analysis is the prediction that the CTD mediates interaction with KorB (Figure 6a) in a way that is very similar to KorA. Modeling of Upf31 with KorB gave a robust ranking_confidence score of 0.64. Removal of the CTD from Upf31 drastically reduced the predicted interaction between KorB and Upf31 to 0.274. When pBP136 KorB was modelled with R751Upf31, which natively lacks a CTD, a similarly low score of 0.252 was predicted. Given the globular domain sequence similarity between the Upf31 of pBP136Km and R751 these results are highly suggestive of pBP136Km Upf31 interacting with KorB via its C-terminus CTD (Figure 6b). Although the Upf31 CTD does not line up with KorA CTD in the way that the CTD of TrbA does, one can detect similarities like the Tyrosine at position 180 and the patch of predominantly basic amino acids. This predicted interaction might allow Upf31 to reduce available KorB either when bound to DNA or free, since we previously observed that KorA and KorB could be copurified in the absence of DNA (McLean et al. 2024). This would in turn explain the increased transcription of the IncP-1 regulatory genes as well as the operons they regulate.

**Figure 6:**
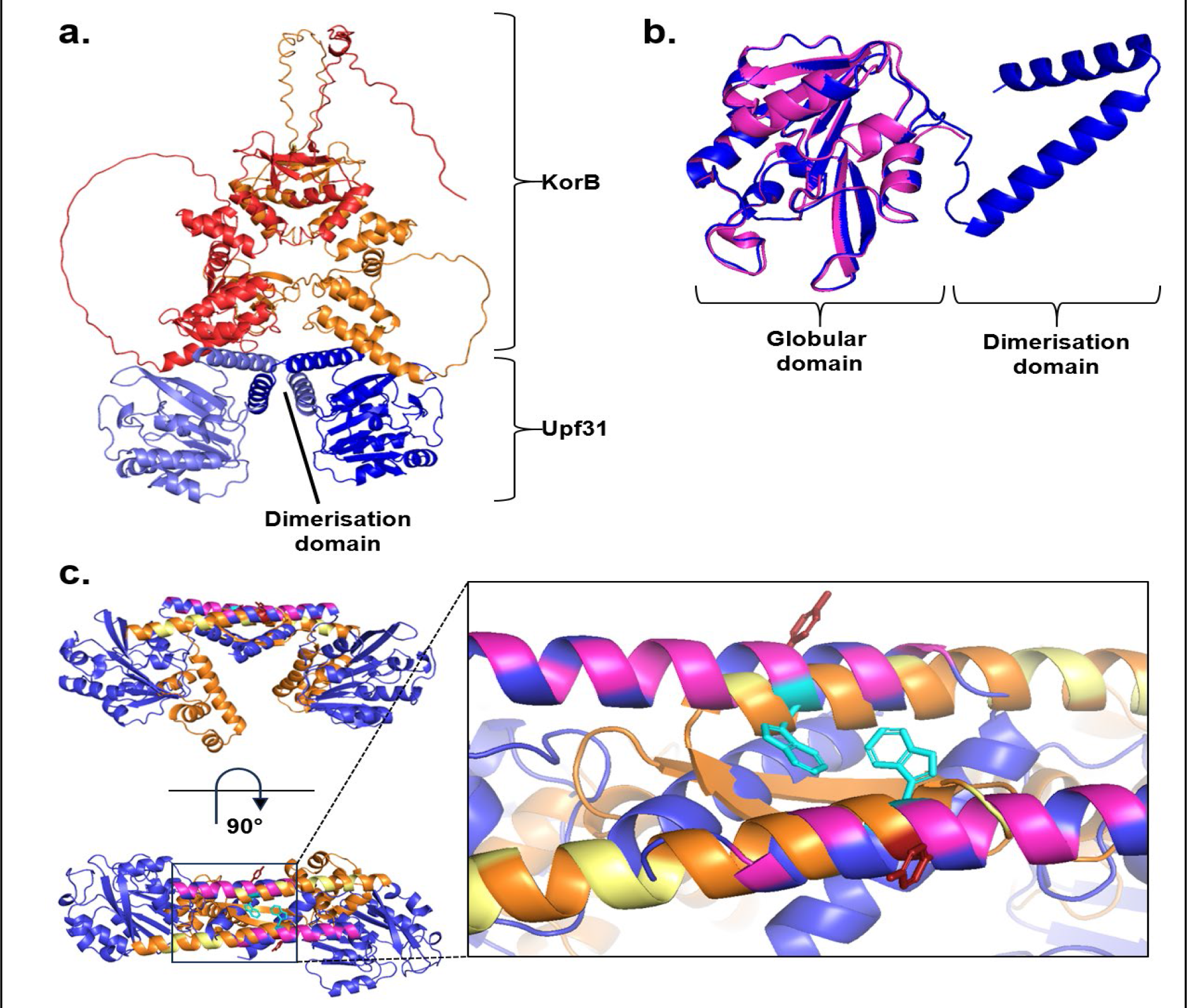
AlphaFold2 modeling reveals the potential interaction between pBP136 Upf31 and KorB is mediated by the C-terminal dimerization domain. **(a)** AlphaFold2 modelling of the pBP136 Upf31 homodimer (light and dark blue) with the pBP136 KorB homodimer (red and orange). **(b)** Superimposed AlphaFold2 models of pBP136 Upf31 (dark blue) and R751 Upf31 (magenta) highlighting the additional C-terminal dimerization domain of pBP136 Upf31. **(c)** Overlay of the AlphaFold2 model of pBP136 Upf31 homodimer (dark blue) with the crystal structure of the RK2 KorA homodimer (PDB: 5CM3, DNA duplex removed). Basic residues in the dimerization domain are highlighted in magenta or yellow. KorA Y84 is highlighted in deep red, Upf31 W198 is highlighted in cyan.

Additional experiments were performed to test the computational prediction of Upf31 requiring a CTD for functionality. Plasmid R751 encodes a truncated Upf31 allele lacking a CTD and persisted very well in *E.* coli but was rapidly lost in the presence of Upf31/pBP136Km expressed *in trans* (Supplemental Figure S2 and Supplemental Data S6). This suggests the CTD of Upf31 is critical to Upf31’s negative effect on plasmid persistence and growth of *E. coli.* To test further whether it is the CTD of pBP136’s Upf31 that is critical for poor plasmid persistence in *E. coli*, rather than other amino acid differences between the two alleles, we tested the effect of the hybrid Upf31 (described earlier) on R751. This hybrid Upf31 combines R751’s truncated Upf31 with the C-terminus of Upf31/pBP136Km and was expressed *in trans* with vector pBR*upf31*. Strikingly, transformation of pBR*upf31* into competent *E. coli* C600 (R751) gave hundreds of pBR*upf31* transformants on L-agar with ampicillin but less than 10 transformants on L-agar with ampicillin and trimethoprim (selecting for both pBR*upf31* and R751). Together our computational and experimental findings strongly suggest that the presence of the CTD of Upf31 from pBP136 is necessary for the poor plasmid persistence phenotype, and likely because of its physical interaction with KorB.

## DISCUSSION

This work sheds new light on the biology and evolution of the largest sampled subgroup of the most intensively studied conjugative plasmids. We demonstrate that the poor persistence in *E. coli* of a conjugative broad-host-range IncP-1β plasmid from *Bordetella pertussis* rapidly improved, even in the absence of selection for the plasmid. The associated drastic amelioration of the plasmid fitness cost to *E. coli* was due to the inactivation of a previously uncharacterized plasmid gene*, upf31,* which affected the expression of the plasmid backbone regulatory circuit including all four major regulators as well as hundreds of chromosomal genes. This change in transcriptome may have been initiated by Upf31 interacting with one of these major plasmid regulators, KorB. Even though plasmids like pBP136 of the incompatibility group IncP-1 are notably persistent in many Proteobacteria (Schmidhauser and Helinski, 1985; De Gelder et al. 2007; Shintani et al. 2010; Yano et al. 2012, 2013, Jain and Srivastava, 2013; Klümper et al. 2015; Kottara et al. 2018;), we show that different alleles of plasmid genes like *upf31* can cause striking differences in plasmid persistence in particular hosts. Understanding how this gene acts and why it may be an advantage in some hosts will expand our understanding of the genetic toolbox plasmids can exploit for their success.

The last decade has seen a flood of seminal studies regarding compensatory evolution of plasmid fitness cost, including multiple studies showing that changes in chromosomal regulatory systems can improve plasmid-host relations. One example is mutations of the *fur* transcriptional regulator involved in iron uptake (Hassan and Troxell, 2013; Stalder et al. 2017), while evolution of the chromosomal *gacA/S* regulators (Harrison et al. 2015) was observed to lower the fitness cost of plasmid carriage within 48 hours of plating on agar (Hall et al. 2020). A few studies have found compensatory evolution via putative chromosomally-encoded helicases (Loftie-Eaton et al. 2017), in one case largely restoring gene expression levels of the plasmid-host pair to that of the plasmid-free bacteria (San Millan et al. 2015). Amid these discoveries, the complex and tightly regulated plasmid transcriptional regulatory systems have perhaps been overlooked as a key factor in plasmid-host relations. One exception was a study which found that a plasmid improved persistence via differential expression of a *parAB* operon, with backbone genes and plasmid regulators otherwise remaining unchanged (Hall et al. 2021). Our work shows that the plasmid encoded Upf31 inactivated during evolution in *E. coli* regulates its own transcription as well as that of the primary plasmid regulators associated with the stable inheritance and control region. Thus, the regulatory systems of *both* plasmids and chromosomes should be included in the growing list of evolutionary pathways that can decrease the fitness cost of a plasmid.

The exact mechanism of action of Upf31 remains experimentally unvalidated but we hypothesize that its C-terminal dimerization domain interacts with plasmid regulator KorB. This was suggested by our use of AlphaFold2 to predict structures and interactions between Upf31 and all other pBP136-encoded proteins, which identified KorB as a highly likely target for Upf31 (Supplemental Data S5). Furthermore, the CTD of Upf31 modeled very similar to the CTD of KorA, which was previously shown to interact with KorB (Bingle et al. 2008). KorB is known to interact co-operatively with the other plasmid backbone repressors (Kostelidou et al. 1999; Kostelidou and Thomas, 2000; Zatyka et al. 2001). Therefore, a KorB-Upf31 interaction that reduces KorB’s ability to interact with other repressors could explain the observed weakening of KorB autogenous control as well as weaker operon repression. These changes in plasmid transcriptional regulation were associated with greatly reduced *E. coli* host fitness and the associated lowered plasmid persistence. It would also align with our result that plasmid R751 with a *upf31* allele that natively lacks a CTD is highly persistent in *E. coli*. In sum, our study suggests that just like KorA, Upf31 may also belong to the family of proteins that can interact with KorB, and that its interaction may override the other interactions with concomitant derepression of gene expression.

The Upf31-associated derepression of global plasmid repressors triggers a secondary effect of large-scale differential gene expression. Thus, the next question becomes which of these differentially expressed genes explains poor *E. coli* fitness. As this negative fitness effect of *upf31* was only observed in the presence of IncP-1β plasmids, we focus here on changes in the plasmid transcriptome. First, of the three plasmids in this study that contained full-length *upf31* homologs, only pALTS29 showed high persistence in *E. coli,* which may suggest low fitness cost. Notably, this plasmid lacked a toxin-antitoxin (TA) system found in nearly all IncP-1β plasmids. This suggests that overexpression of TA genes may explain poor *E. coli* fitness.

However, the higher persistence of pALTS29 could equally be explained by differences in Upf31 amino acid sequences (Supplemental Table S2). Given our hybrid Upf31 results, there are only two amino acid differences between the Upf31/R751/pBP136Km hybrid and Upf31/pALTS29 that could explain differences in plasmid persistence, i.e. at positions 82 and 92. Another possible explanation for the negative effect of Upf31 on *E. coli* fitness is a set of four genes in the control region that were upregulated in the presence of Upf31 (*kleA, kleE, klcA, klcB,* or ‘*kil’ genes*). These genes have been shown to have harmful effects on the growth of *E.coli* strains when not repressed by the global Kor regulators (Figurski et al. 1982; Kornacki et al. 1993; Larsen and Figurski, 1994), with *klcB* found harmful only in the presence of an IncP-1 plasmid (Bhattacharyya and Figurski, 2001). Both above hypotheses assume that the observed mis- regulation of *non-*regulatory plasmid genes cause physiological change within the bacterial host which would explain the responsive shift in chromosomal gene expression levels. An alternative model is that the observed mis-regulation of the plasmid repressors (KorA/B/C) also directly changes expression of chromosomal genes in an example of plasmid-chromosome ‘cross-talk’ (Vial and Hommais, 2020; Hall et al. 2021; Marincola et al. 2021; Thompson et al. 2023). Future studies will have to identify the specific overexpressed plasmid genes that have such a negative effect on *E. coli* fitness and by what mechanism, and if the differentially transcribed chromosomal genes are at all responsible. It also remains to be determined whether *upf31* provides an advantage in the original *B. pertussis* host.

Upf31 might be an important clue to understanding the four groups of conserved inverted repeats (IR) that were previously described (Thorsted et al. 1998) in IncP-1b plasmids but without any explanation of purpose or function. The *upf31* gene itself is embedded in one of the four IR regions and appears to autoregulate its own expression by binding to one of the IR regions. It is therefore likely that Upf31 binds to all the IR sequences although we currently have no indication of purpose. However, it is noteworthy that *klcB* is among the highest overexpressed genes in the presence of Upf31 (Figure 6) and includes several IR within its coding sequence which may correlate with being available to compete with KorB for repression.

We next wish to report two important lessons we learned during this study. First, we frequently observed rapid loss of *upf31* during routine laboratory cultivation due to various deletions, prompting us to always sequence the region of *upf31* to confirm its presence. This suggests that rapid evolution of costly plasmid genes may often go unrecognized during cultivation and plasmid transfer between strains by matings and electroporation. Our work aligns with previous findings of rapid (<48 hours) compensatory evolution between plasmid-host pairs, even without known selection (Hall et al. 2020). Second, our work showed that Upf31 is likely not a functional methylase in contrast to automated annotations describing the gene as such (without experimental data) in many plasmid genomes. This highlights the argument that plasmid genes *without* experimental data on their function should be assigned locus tags rather than putative function-related gene names (Thomas et al. 2017).

In conclusion, despite decades of great progress in plasmid biology and the pivotal role of conjugative plasmids in antibiotic resistance dissemination, a substantial portion of plasmids harbor multiple genes with unknown functions. Our study highlights the importance of one such gene, which when inactivated led to a lasting host-plasmid pair, effectively rescuing a plasmid from potential extinction (Gomulkiewicz and Holt, 1995) in a bacterial population. The investigation of this specific gene revealed that this evolutionary rescue is closely linked to alterations in the plasmids regulatory circuit. These findings underscore the significance of identifying and experimentally validating uncharacterized plasmid genes to understand how evolution underwrites the spread and persistence of plasmids in bacterial communities.

## MATERIALS AND METHODS

### Bacterial strains, plasmids, and media

IncP-1β plasmid pBP136 was discovered in a strain of *Bordetella pertussis* isolated from a lethal case of infant whooping cough^35^. A kanamycin gene was later inserted to generate pBP136Km (NCBI accession number NZ_OR146256.1), which provided a selectable marker in the originally cryptic plasmid^56^. The expression vector pCW-LIC-*upf31* was derived from pCW- LIC-*sacB* as described below in the cloning section.

*Escherichia coli* K-12 MG1655 was derived^57^ from an isolate within the stool of a diphtheria patient in 1925, and *E. coli* JM109 and DH5α^58^ are later derivatives of this strain with useful attributes relevant to cloning. We generated rifampicin and nalidixic resistant clones of *E. coli* K-12 MG1655 by recovering resistant mutants after plating on agar with the respective antibiotic. New constructs and initial, intermediate, and final populations of the plasmid persistence assays were all archived at -70 °C in 30% glycerol.

All bacteria in this study were grown at 37°C in lysogeny broth (LB) shaken at 200 RPM or on LB agar (LBA) plates. Strains were grown with 50 μg ml^-1^ kanamycin (km) or 100 μg ml^-1^ ampicillin (amp) when appropriate for plasmid maintenance. Expression vectors were induced with 500 μM isopropyl β- d-1-thiogalactopyranoside (IPTG) when induction is mentioned in the text.

### Plasmid Persistence Assay

All plasmid persistence assays were performed in triplicate and first grown overnight (O/N) with kanamycin to select for initial plasmid maintenance (time T0). Daily serial transfers of 4.9 μL of overnight culture into 5 mL of sterile broth *without* antibiotics were then performed for ten more days (times T1-T10). Each day the cultures were diluted 10^-6^ in 1x phosphate buffered saline (PBS) and 100 uL was spread onto dilution plates. The daily plasmid-containing fraction of the population was determined by replica plating 52 randomly chosen colonies from the dilution plates onto LB plates with and without kanamycin.

### Growth Assays

We performed growth assays in batch culture to measure the effect of Upf31 on growth rates and final densities. Strains were grown from freezer archives for 24 hours and diluted 1:100, then grown another 24 hours and finally diluted 1:1000 in sterile media. From these ten biological replicates each were loaded at a volume of 200uL into the respective wells of a 96- well plate. A SPECTROstar Nano plate reader measured optical density at 600nm every ten minutes for 23 hours with incubation at 37C and 500 RPM orbital shaking. The R package GrowthRates (Hall et al. 2014) v. 0.8.4 was used to calculate maximum growth rate using default parameters.

### Low-Density Competition Assays

The low-density competition assay was developed in this study to avoid plasmid transfer during the assay. It contains two distinct steps: (*i*) determining the likely “conjugation-free” time window for specific low donor and recipient densities, and (*ii*) performing a traditional competition assay at low densities within the established time window.

To find the likely window of time before density is high enough for conjugation to occur, we used K-12 MG1655 strains^34^ with spontaneous mutations conferring resistance to nalidixic acid (Nal) and rifampicin (Rif) (donor K-12N(pBP136Km) and recipient K-12R, respectively). Donor and recipient were mixed1:1 at densities of 10^2^/ml into 2.5 mL LB shaken 200 RPM at 37°C. To select for possible transconjugants, the cultures were mixed hourly with another 2.5 mL of LB with Km and Rif to create a 5mL mixture with 50 μg ml^-1^ Km and 50 μg ml^-1^ Rif.

These were grown 24 hours after which the earliest timepoint showing turbidity was associated with the emergence of transconjugants K-12R(pBP136Km). The first turbid culture emerged after six hours, and therefore a five-hour window of likely conjugation-free growth was chosen for the subsequent competition assays.

Competition assays were then performed using K-12 vs. K-12(pBP136Km) and K-12 vs. K-12(pBP136KmΔ*upf31*). Each experiment began with 1:1 mixtures at densities of 10^2^/ml in 5 LB without antibiotics in shaken test tubes. Plating onto LBA occurred at the start time and after five hours (after appropriate dilutions), and the relative increase in cell numbers (cfu/ml) was used as a proxy for relative fitness as follows: (Wiser and Lenski, 2015)

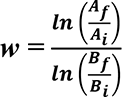

where *w* is the Malthusian relative fitness term, *AI* and *AF* are the initial and final plasmid containing K-12 populations, respectively, and *BI* and *BF* are the initial and final plasmid-free K- 12 populations, respectively.

### Methylation Detection

To test if *upf1* encodes a methyltransferase as computationally predicted we utilized the base modification analysis provided by the Pacbio Sequel II DNA sequencer. The methylase-free *E. coli* strain ER2796^43^ with expression vector pCW-LIC-*upf31* was tested against ER2796 (pCW-LIC-*sacB*) for differential base modification^36^ on a Pacbio Sequel II at the Arizona Genomic Institute in Tucson, AZ, USA. Both strains were grown with antibiotic selection and 500 μM IPTG induction before genomic extraction using the Sigma GeneElute Bacterial kit. Base Modification Analysis was run on SMRT Link v. 10.2.0.133434 with a reference made by merging the *E. coli* assembly with the respective plasmids and enabling “Find Modified Base Motifs” and “Consolidate Mapped BAMs for IGV” options. All the other options were left as default.

### RNA-Seq

To understand how *upf31* and pBP136Km interact to influence *E. coli* fitness, we performed RNA-seq. Strains K-12WT (pBP136Km) and K-12WT (pBP136KmΔ*upf31)* were grown overnight in 5 ml LB with Km and centrifuged. The cell pellets were sent to Zymo Research for Total RNA-Seq Service. Sequencing libraries were constructed from total RNA samples and were prepared using the Zymo-Seq RiboFree Total RNA Library Prep Kit (Cat # R3000) according to the manufacturer’s instructions (https://www.zymoresearch.com/products/zymo-seq-ribofree-total-rna-library-kit). RNA-Seq libraries were sequenced on an Illumina NovaSeq to a sequencing depth of at least 30 million read pairs (150 base paired-end sequencing) per sample.

A custom bioinformatics pipeline began with TrimGalore! v. 0.67 to remove adapters and low-quality reads. Bowtie2 (Langmead and Salzberg, 2012) v. 2.4.5 then mapped reads to merged chromosomal and plasmid references. These were counted using featureCounts(Liao et al. 2014) v. 2.0.1, the output of which was input unto DESeq2 (Love et al. 2014) v. 1.38.1, for final calculations of relative expression.

### DNA Sequencing

Whole genome sequencing was performed by SeqCenter, LLC in Pittsburg, PA, USA. Genomic DNA extractions were performed using Sigma GeneElute Bacterial kit. The libraries were prepared using the Illumina DNA Prep kit with IDT 10bp UDI indices and sequenced on an Illumina NextSeq 2000 with 2x151bp reads. Demultiplexing, quality control and adapter trimming was initially performed using Illumina’s bcl-convert v. 3.9.3. TrimGalore! v. 0.6 was additionally used to remove remaining adapters and low-quality reads. Breseq (Barrick et al. 2014) v. 0.36.0 was used to identify mutations between ancestral and evolved strains.

### Cloning

pCW-LIC (hereafter pCW-LIC-*sacB*) was a gift from Cheryl Arrowsmith (Addgene plasmid #26098; http://n2t.net/addgene:26098; RRID:Addgene_26098). Plasmid pCW-LIC-*sacB* was double cut at the *NdeI* and *HindIII* restrictions sites, which created a 4,962 base pair (bp) linearized backbone and a 2,302 bp linear fragment containing *sacB* and its promoter. These were separated and the linearized backbone recovered from an agarose gel using a Thermo Scientific (TS) GeneJET Gel Extraction Kit. Next, gene *upf31* was amplified from pBP136Km miniprep using upstream primer 5’-*GGTGGT*<u>CATATG</u>TCCAGGAAGAAGGCCATGAG-3’ and downstream primer 5’- *GGTGG*<u>TAAGCTT</u>CTACTCGGCCGCCTCTAG-3’ (flanking sequences for restriction enzymes in italics, *NheI* and *HindIII* sites underlined, respectively). The ends of the *upf31* amplicon were then double digested at the *NdeI* and *HindIII* sites, and the small digested ends removed using a TS GeneJET PCR Purification Kit. The *upf31* segment and linearized pCW-LIC backbone were joined using T4 DNA ligase and the now-circularized pCW- LIC-*upf31* was electroporated into DH5α using standard methods(Sambrook and Russell, 2001).

To construct the upf31p-*xylE* reporter gene fusion complementary 68-nt oligomers were designed to create a double stranded fragment with *Xba*I and *Eco*RI sticky ends for insertion into *xylE* reporter plasmid pGCMT1 to create pGCMT-upf31p as shown in Supplementary Figure S6. The 68 bp region upstream of *upf31* was from plasmid R751.

To construct the R751-pBP136 hybrid *upf31*, translational start plus codons 1 to 173 were amplified by PCR with primers 5’-TGCAAGCTTTAATGCGGTAGCCAAGTCCCGATTTACTCCAG-3’ and 5’- CCGGCTCGGGGTAGTTCATC-3’ from R751 template. To create codons 174 to 224 from pBP136 PCR-driven overlapping(Heckman and Pease, 2007) of synthetic oligomers (104 nt and 117 nt) were utilized with pBP136 template. The whole segment was amplified with primers 5’- TGCAAGCTTTAATGCGGTAGCCAAG-3’ and 5’-TCGGTCGACGCAGGCGTGAC-3’ and after cutting with restriction enzymes inserted into both pBR322 (Accession J01749.1) and pACYC184 (Accession X06403.1) between their HindIII and SalI sites so that *upf31* was transcribed from the *tetA* promoter.

### *xylE* reporter assay

Overnight LB cultures were inoculated from single colonies of strains with reporter plasmid pGCMT1 or its derivative pGCMTupf31p alone (with just kanamycin selection) or with a second plasmid vector (pBR322) or vector plus the hybrid *upf31* gene (with both kanamycin and ampicillin) and incubated for 16 h. Bacteria from 1 ml aliquots were pelleted, resuspended in 0.5 ml sonication buffer, sonicated with three bursts of 5 seconds, the cell debris cleared with 10 min centrifugation at maximum speed in a microfuge at 4°C and then measured quantitatively for XylE activity and protein concentration as described previously (Zukowski et al. 1983).

### Upf31 protein expression

Induction cultures of pCW-LIC-upf31 in *E. coli* K-12 M1655 were started from an overnight liquid culture of LB (Lennox) with 100 µg/ml ampicillin (LB Amp). Twenty-five milliliter LB Amp cultures were initiated at OD600 0.05 using the liquid overnight culture to serve as the induction cultures. The 25 milliliter cultures were grown at 37° C while shaking at 200 rpm. When the 25 milliliter cultures reached OD600 > 1.0, one milliliter of culture was removed, and the cells were harvested by centrifugation for four minutes at 16,000 x g at room temperature in an Eppendorf 5425 centrifuge. This material served as the pre-induction (or 0 hour) time point for each culture. ITPG was then added to each induction culture to a final concentration of 0.1 mM, 0.5 mM, or 1 mM and growth was monitored at OD600. At the time points indicated in Figure S2, one milliliter of culture was removed from each culture and cells were isolated as above. Cells were resuspended in 1x Tris-Glycine running buffer (25 mM Tris-Cl, pH 8.3, 192 mM glycine, and 0.1% SDS) at 100 microliters per 1.0 = OD600 cells. Laemmli buffer from Bio-Rad was added to the resuspended cell pellets to a final concentration of 1x and heated 10 minutes at > 98° C prior to gel loading.

### Upf31 SDS-Page Gel Electrophoresis

Heated, denatured cellular extract (see previous section) samples were loaded on a Tris-glycine gel with a 12% acrylamide resolving gel and a 4% acrylamide stacking gel. Each sample lane contained 0.1 OD600 of cells. The molecular weight standards used were unstained broad range protein standards (NEB) and then the gel was run at 100 volts until the dye front was 1 cm from the bottom of the gel and then stained one hour in Coomassie blue stain (50% methanol, 10% acetic acid, 0.25% Coomassie brilliant blue R-250) followed by destaining (5% methanol, 7% acetic acid) overnight.

### Computational Modeling of Interactions between Upf31 and pBP136-encoded Proteins

An ‘*in silico* pulldown’ of Upf31 was carried out using the LazyAF pipeline(McLean, 2024). In short, the pBP136 genome (AB237782) was retrieved from NCBI GenBank as a FASTA protein coding sequence. The Upf31 protein coding sequence was used to generate individual concatenated FASTA files with Upf31 and each pBP136 coding sequence. ColabFold v1.5.5: AlphaFold2 w/ MMseqs2 BATCH was then run on Google Colaboratory using the High-RAM A100 GPU with the following settings: msa_mode: MMseqs2 (UniRef+Environmental), num_models: 5, num_recycles: 3, stop_at_score: 100. Subsequently the JSON files were analyzed to retrieve the pTM and ipTM scores for each top ranked model and calculated the ranking_confidence score (0.2 pTM + 0.8 ipTM).

## ACKNOWLEDGEMENTS

We acknowledge Tùng Lê at the John Innes Centre (UK) for his work on the *in silico* Upf31 pulldown. Dr. Ben Kerr at the University of Washington helped O.K design the low-density competition assay. We are indebted for the laboratory efforts of undergraduate researchers Luke Hoover, Morgan Sower, Sue Winger, Katlyn Schafer, Courtney Stattner, Mattie Hagestad, Audrey Dingel, and Alexandra Gal and research technician Jack Millstein at the University of Idaho. We also thank Salvador (“Chava”) Castaneda Barba for improving this manuscript. The project was made possible with the skilled sequencing efforts of IIDS Genomics and Bioinformatics Resources Core (GBRC) at the University of Idaho as well as the Arizona Genomics Institute at the University of Arizona.

## FUNDING RESOURCES

E.M.T.and C.E. received partial support for this work from the National Institute of Allergy and Infectious Diseases Extramural Activities grant no. R01AI084918 from the National Institutes of Health. C.A.E. was also supported by the National Science Foundation Graduate Research Fellowship grant no. DGE-2019265372 as well as the Bioinformatics and Computational Biology (BCB) Fellowship and the Paul Joyce Memorial BCB Fellowship Endowment at the University of Idaho. O.K. was supported by the National Science Foundation Graduate Research Fellowship grant no. DGE-1762114. This work is supported by the Wellcome Trust Investigator grant 221776/Z/2/Z to Tùng Lê that supported T.C.M, and by the BBSRC funded Institute Strategic Program Harnessing Biosynthesis for Sustainable Food and Health (HBio) (BB/X01097X/1). The funders had no role in study design, data collection and analysis, decision to publish, or preparation of the manuscript.

