## Supplemental Data S2 for "Evolution of a Plasmid Regulatory Circuit Ameliorates Plasmid Fitness Cost"

Phyre Home | Retrieve Phyre Job Id | Fetch

Phyre2

| | |
|---|---|
| Email | |
| Description | upf31_pBP136Km |
| Date | Thu Apr 29 18:42:12 BST 2021 |
| Unique Job ID | 404c522ea6d0d006 |
| Sequence | MRYPGGKGGA ... Download FASTA |
| Job Type | **normal** |
| Job Expiry | |

Download zip of all results

#### Summary

Top model

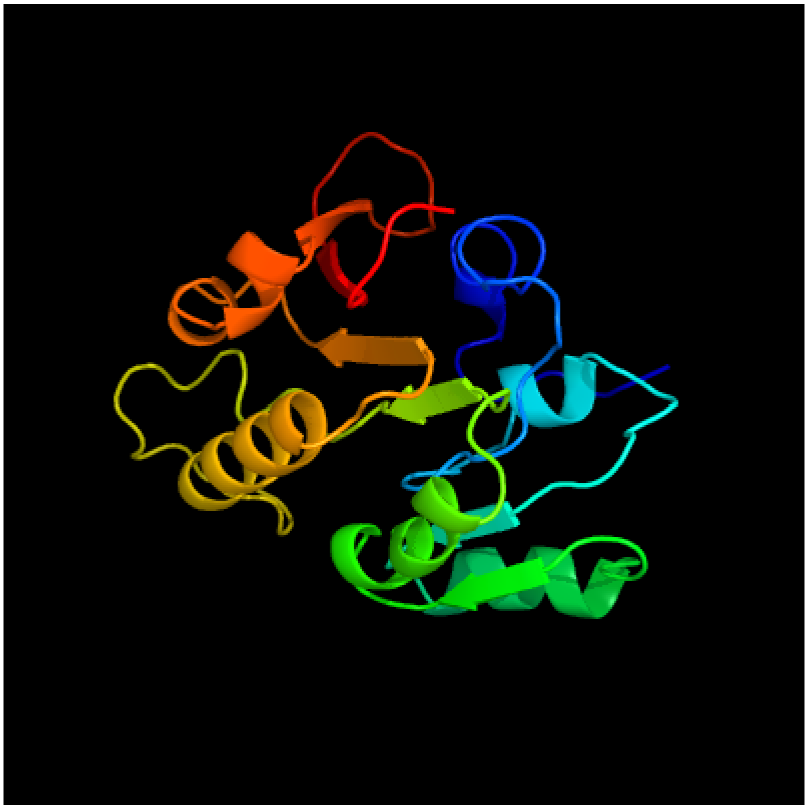

Image coloured by rainbow N → C terminus

Model dimensions (Å): **X**:45.259 **Y**:38.600 **Z**:41.472

Model (left) based on template d2dpma_

Top template information

**Fold:**S-adenosyl-L-methionine-dependent methyltransferases
**Superfamily:**S-adenosyl-L-methionine-dependent methyltransferases
**Family:**N6 adenine-specific DNA methylase, DAM

Confidence and coverage

Confidence: 100.0% Coverage: 70%

157 residues ( 70% of your sequence) have been modelled with 100.0% confidence by the single highest scoring template.

3D viewing

Interactive 3D view in JSmol

For other options to view your downloaded structure offline see the FAQ

#### Sequence analysis

View PSI-Blast Pseudo-Multiple Sequence Alignment | Download FASTA version

Secondary structure and disorder prediction [Show]

Domain analysis [Show]

Detailed template information [Show]

Binding site prediction

Automated 3DLigandsite submission is temporarily suspended due to server load. However you may submit models directly to 3DLigandSite HERE

---

Phyre is now FREE for commercial users!

All images and data generated by Phyre2 are free to use in any publication with acknowledgement

**Please cite:** The Phyre2 web portal for protein modeling, prediction and analysis.

Kelley LA *et al.*. *Nature Protocols* 10, 845-858 (2015)[pdf] [Citation link]

If you use the binding site predictions from 3DLigandSite, **please also cite:**

3DLigandSite: predicting ligand-binding sites using similar structures.

Wass MN, Kelley LA and Sternberg MJ *Nucleic Acids Research* 38, W469-73 (2010) [PubMed]

© Structural Bioinformatics Group
Imperial College London

Lawrence Kelley, Michael Sternberg

Disclaimer

Terms and Conditions

**Component software**

Template detection: HHpred 1.51

Secondary structure prediction: Psi-pred 2.5

Disorder prediction: Disopred 2.4

Transmembrane prediction: Memsat_SVM

Multi-template modelling and *ab initio*: Poing 1.0

Imperial College London

| | |
|---|---|
| Email | |
| Description | upf31_pBP136Km |
| Date | Thu Apr 29 18:42:12 BST 2021 |
| Unique Job ID | 404c522ea6d0d006 |

#### Secondary structure and disorder prediction

| | 1–60 |
|---|---|
| Sequence | MRYPGGKGGAGVYQTIINNIPPHDTYIETHLGGGNILERKRPAARSIGIDIDPEVIEVWR |
| Disorder | ? ? ? ? ? ? ? ? ? ? ? ? ? ? ? ? |

| | 61–120 |
|---|---|
| Sequence | QLDVPGLELYHGDAVDWLESHQFTGSEFVYADPPYVMDSRRGGKLYRYEYDDADHVRLLD |
| Disorder | ? ? ? ? ? ? ? |

| | 121–180 |
|---|---|
| Sequence | VLARLPCAVMVSGYSSTIYDSSPLASWRTIDFNAMTRGGIAIERLWMNYPEPAKLHDLRY |
| Disorder | ? ? ? ? ? ? ? ? ? |

| | 181–224 |
|---|---|
| Sequence | LGSNFRERERIKRKKARWQAKIAKLDPLERAAIMECLRELEAAE |
| Disorder | ? ? ? ? ? ? ? ? ? ? ? ? ? ? ? |

Confidence Key

High(9) … Low (0)

? Disordered ( 21%)

Alpha helix ( 44%)

Beta strand ( 14%)

### Phyre2

| | |
|---|---|
| Email | |
| Description | upf31_pBP136Km |
| Date | Thu Apr 29 18:42:12 BST 2021 |
| Unique Job ID | 404c522ea6d0d006 |

#### Domain analysis

| Rank | Aligned region |
|---|---|
| 1 | d2dpma_ |
| 2 | d1yf3a1 |
| 3 | c2g1pB_ |
| 4 | c7m6bB_ |
| 5 | d2fpoa1 |
| 6 | c3lduA_ |
| 7 | c3ldgA_ |
| 8 | c3egiA_ |
| 9 | c3v8vB_ |
| 10 | c3vseA_ |
| 11 | d2esra1 |
| 12 | c5e72A_ |
| 13 | c3p9nA_ |
| 14 | d2as0a2 |
| 15 | d2b78a2 |
| 16 | c1wxwA_ |
| 17 | d1wxxa2 |
| 18 | c3k0bA_ |
| 19 | c3tm4A_ |
| 20 | c2as0A_ |
| 21 | |
| 22 | |
| 23 | |
| 24 | |
| 25 | |
| 26 | |
| 27 | |
| 28 | |
| 29 | |
| 30 | |
| 31 | |
| 32 | |
| 33 | |
| 34 | |
| 35 | |
| 36 | |
| 37 | |
| 38 | |
| 39 | |
| 40 | |
| 41 | |
| 42 | |
| 43 | |
| 44 | |
| 45 | |
| 46 | |
| 47 | |
| 48 | |
| 49 | |
| 50 | |
| 51 | |
| 52 | |
| 53 | |
| 54 | |
| 55 | |
| 56 | |
| 57 | |
| 58 | |

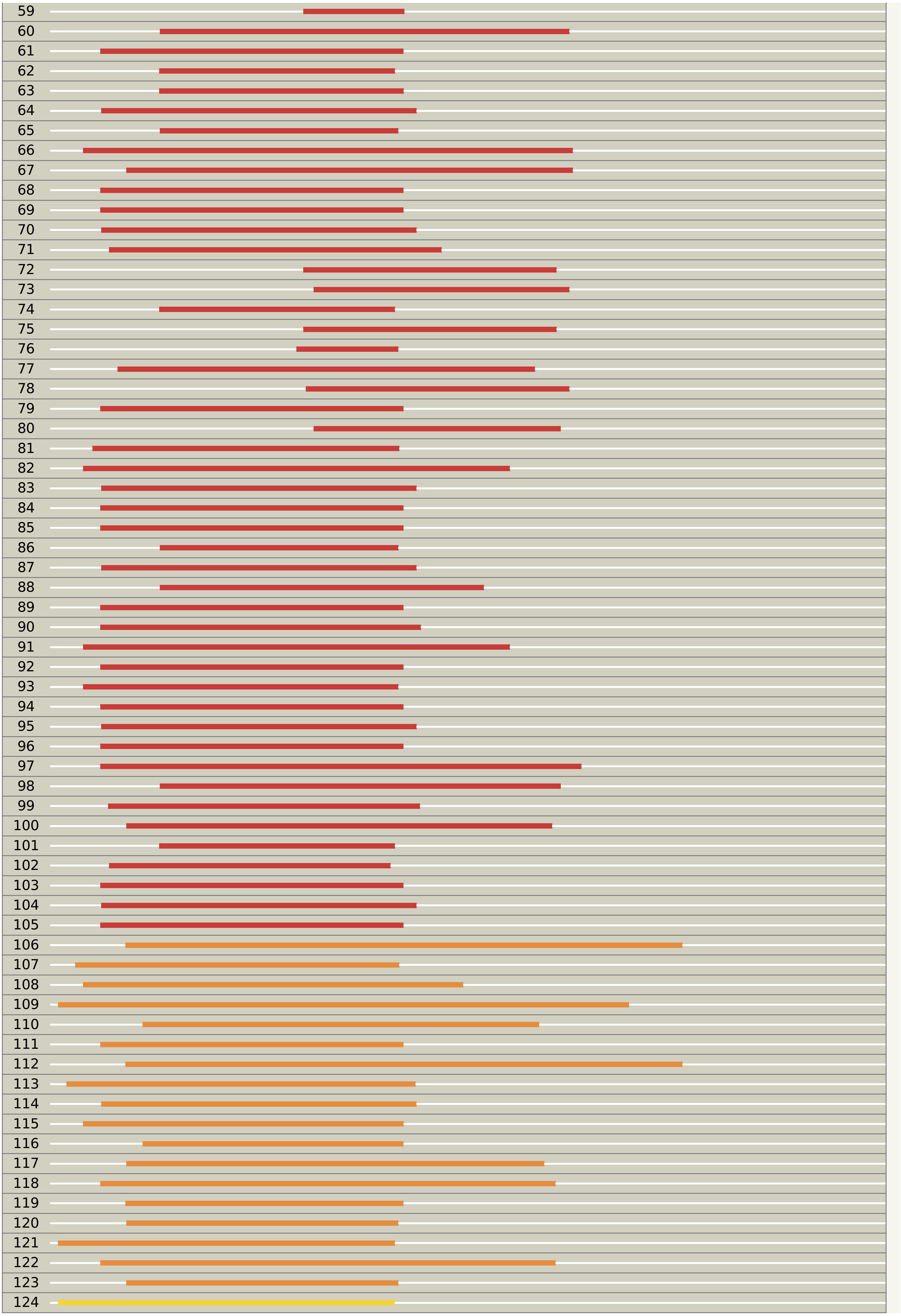

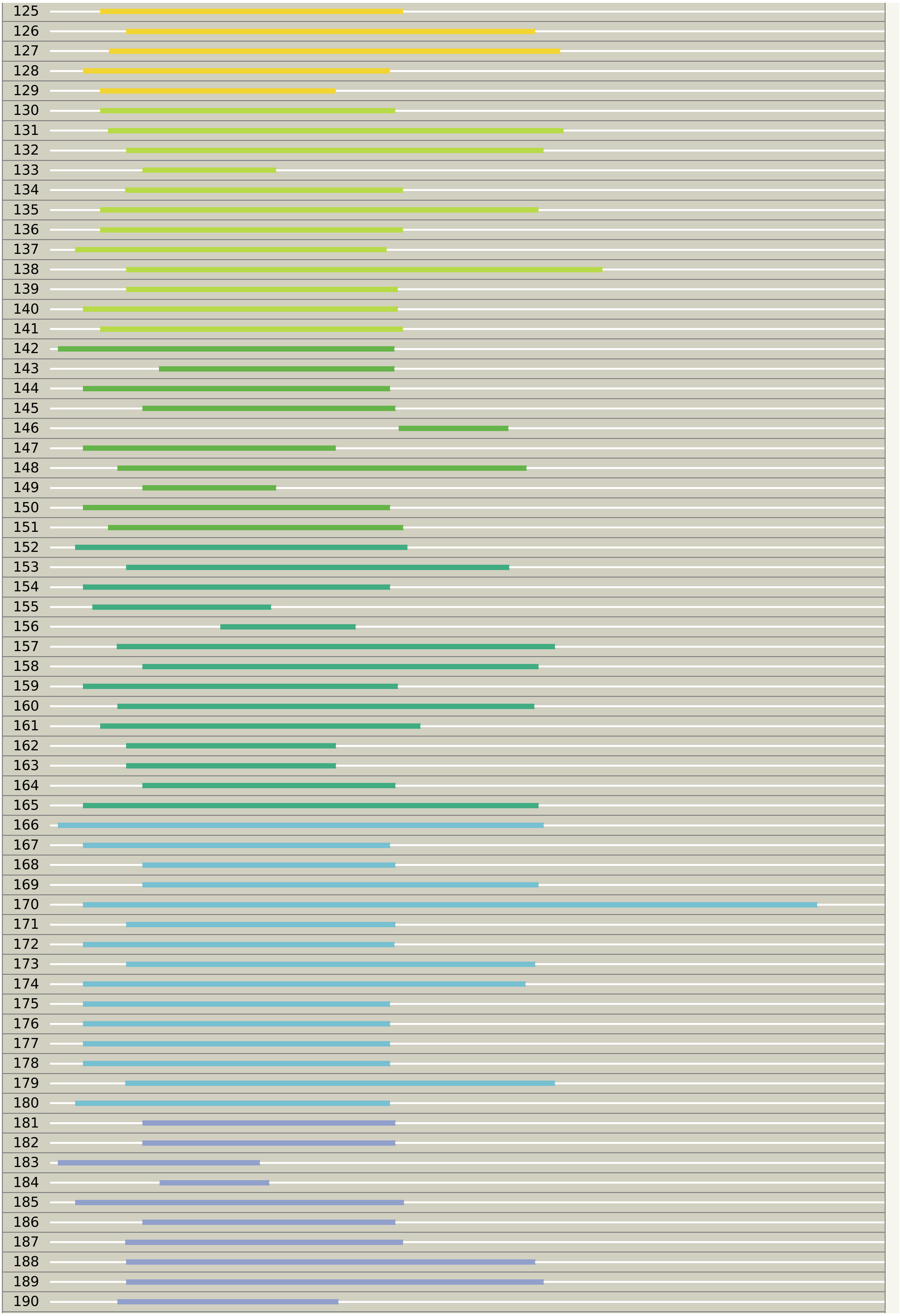

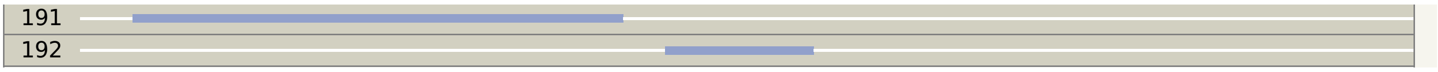
191
192

Phyre²

| | |
|---|---|
| Email | |
| Description | upf31_pBP136Km |
| Date | Thu Apr 29 18:42:12 BST 2021 |
| Unique Job ID | 404c522ea6d0d006 |

Detailed template information

| # | Template | Alignment Coverage | 3D Model | Confidence | % i.d. | Template Information |
|---|---|---|---|---|---|---|
| 1 | d2dpma_ | Alignment | 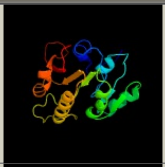 | 100.0 | 29 | **Fold:**S-adenosyl-L-methionine-dependent methyltransferases **Superfamily:**S-adenosyl-L-methionine-dependent methyltransferases **Family:**N6 adenine-specific DNA methylase, DAM |
| 2 | d1yf3a1 | Alignment | 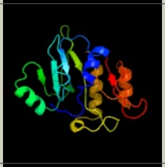 | 100.0 | 22 | **Fold:**S-adenosyl-L-methionine-dependent methyltransferases **Superfamily:**S-adenosyl-L-methionine-dependent methyltransferases **Family:**N6 adenine-specific DNA methylase, DAM |
| 3 | c2g1pB_ | Alignment | 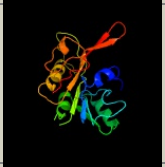 | 100.0 | 22 | **PDB header:**transferase/dna **Chain:** B: **PDB Molecule:**dna adenine methylase; **PDBTitle:** structure of e. coli dna adenine methyltransferase (dam) |
| 4 | c7m6bB_ | Alignment | 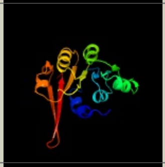 | 100.0 | 34 | **PDB header:**transferase **Chain:** B: **PDB Molecule:**site-specific dna-methyltransferase (adenine-specific); **PDBTitle:** the crystal structure of mcbe1 |
| 5 | d2fpoa1 | Alignment | 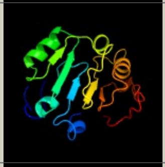 | 98.1 | 21 | **Fold:**S-adenosyl-L-methionine-dependent methyltransferases **Superfamily:**S-adenosyl-L-methionine-dependent methyltransferases **Family:**YhhF-like |
| 6 | c3lduA_ | Alignment | 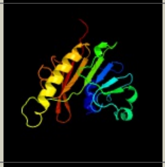 | 98.1 | 20 | **PDB header:**transferase **Chain:** A: **PDB Molecule:**putative methylase; **PDBTitle:** the crystal structure of a possible methylase from2 clostridium difficile 630. |
| 7 | c3ldgA_ | Alignment | 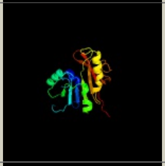 | 98.0 | 18 | **PDB header:**transferase **Chain:** A: **PDB Molecule:**putative uncharacterized protein smu.472; **PDBTitle:** crystal structure of smu.472, a putative methyltransferase complexed2 with sah |
| 8 | c3egiA_ | Alignment | 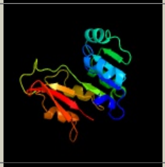 | 98.0 | 14 | **PDB header:**transferase **Chain:** A: **PDB Molecule:**trimethylguanosine synthase homolog; **PDBTitle:** methyltransferase domain of human trimethylguanosine synthase tgs12 bound to m7gpppa (inactive form) |
| 9 | c3v8vB_ | Alignment | 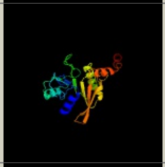 | 97.9 | 20 | **PDB header:**transferase **Chain:** B: **PDB Molecule:**ribosomal rna large subunit methyltransferase l; **PDBTitle:** crystal structure of bifunctional methyltransferase ycby (rlmlk) from2 escherichia coli, sam binding |
| 10 | c3vseA_ | Alignment | 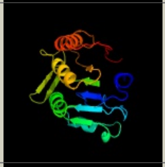 | 97.9 | 13 | **PDB header:**transferase **Chain:** A: **PDB Molecule:**putative uncharacterized protein; **PDBTitle:** crystal structure of methyltransferase |
| 11 | d2esra1 | Alignment | 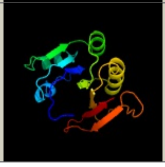 | 97.9 | 15 | **Fold:**S-adenosyl-L-methionine-dependent methyltransferases **Superfamily:**S-adenosyl-L-methionine-dependent methyltransferases **Family:**YhhF-like |

| | | | | | | |
|---|---|---|---|---|---|---|
| 12 | c5e72A_ | Alignment | 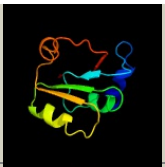 | 97.8 | 26 | **PDB header:**transferase<br>**Chain:** A: **PDB Molecule:**n2, n2-dimethylguanosine trna methyltransferase;<br>**PDBTitle:** crystal structure of the archaeal trna m2g/m22g10 methyltransferase2 (atrm11) in complex with s-adenosyl-l-methionine (sam) from3 thermococcus kodakarensis |
| 13 | c3p9nA_ | Alignment | 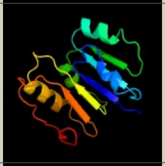 | 97.7 | 22 | **PDB header:**transferase<br>**Chain:** A: **PDB Molecule:**possible methyltransferase (methylase);<br>**PDBTitle:** rv2966c of m. tuberculosis is a rsmd-like methyltransferase |
| 14 | d2as0a2 | Alignment | 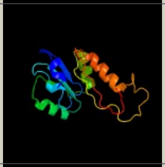 | 97.7 | 21 | **Fold:**S-adenosyl-L-methionine-dependent methyltransferases<br>**Superfamily:**S-adenosyl-L-methionine-dependent methyltransferases<br>**Family:**hypothetical RNA methyltransferase |
| 15 | d2b78a2 | Alignment | 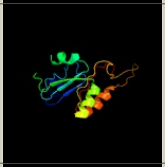 | 97.7 | 15 | **Fold:**S-adenosyl-L-methionine-dependent methyltransferases<br>**Superfamily:**S-adenosyl-L-methionine-dependent methyltransferases<br>**Family:**hypothetical RNA methyltransferase |
| 16 | c1wxwA_ | Alignment | 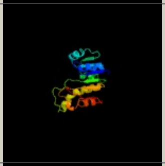 | 97.6 | 11 | **PDB header:**transferase<br>**Chain:** A: **PDB Molecule:**hypothetical protein ttha1280;<br>**PDBTitle:** crystal structure of tt1595, a putative sam-dependent2 methyltransferase from thermus thermophillus hb8 |
| 17 | d1wxxa2 | Alignment | 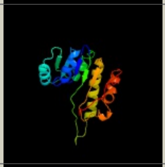 | 97.6 | 11 | **Fold:**S-adenosyl-L-methionine-dependent methyltransferases<br>**Superfamily:**S-adenosyl-L-methionine-dependent methyltransferases<br>**Family:**hypothetical RNA methyltransferase |
| 18 | c3k0bA_ | Alignment | 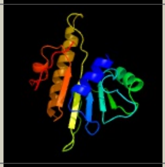 | 97.6 | 24 | **PDB header:**structural genomics, unknown function<br>**Chain:** A: **PDB Molecule:**predicted n6-adenine-specific dna methylase;<br>**PDBTitle:** crystal structure of a predicted n6-adenine-specific dna methylase2 from listeria monocytogenes str. 4b f2365 |
| 19 | c3tm4A_ | Alignment | 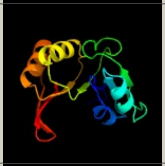 | 97.5 | 18 | **PDB header:**transferase<br>**Chain:** A: **PDB Molecule:**trna (guanine n2-)-methyltransferase trm14;<br>**PDBTitle:** crystal structure of trm14 from pyrococcus furiosus in complex with s-2 adenosylmethionine |
| 20 | c2as0A_ | Alignment | 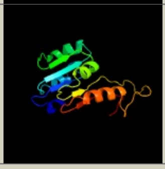 | 97.4 | 20 | **PDB header:**transferase<br>**Chain:** A: **PDB Molecule:**hypothetical protein ph1915;<br>**PDBTitle:** crystal structure of ph1915 (apc 5817): a hypothetical rna2 methyltransferase |
| 21 | d2fhpa1 | Alignment | not modelled | 97.4 | 23 | **Fold:**S-adenosyl-L-methionine-dependent methyltransferases<br>**Superfamily:**S-adenosyl-L-methionine-dependent methyltransferases<br>**Family:**YhhF-like |
| 22 | c3c0kB_ | Alignment | not modelled | 97.4 | 15 | **PDB header:**transferase<br>**Chain:** B: **PDB Molecule:**upf0064 protein yccw;<br>**PDBTitle:** crystal structure of a ribosomal rna methyltranferase |
| 23 | d1ws6a1 | Alignment | not modelled | 97.4 | 13 | **Fold:**S-adenosyl-L-methionine-dependent methyltransferases<br>**Superfamily:**S-adenosyl-L-methionine-dependent methyltransferases<br>**Family:**YhhF-like |
| 24 | c4dmgA_ | Alignment | not modelled | 97.4 | 19 | **PDB header:**transferase<br>**Chain:** A: **PDB Molecule:**putative uncharacterized protein ttha1493;<br>**PDBTitle:** thermus thermophilus m5c1942 methyltransferase rlmo |
| 25 | c2b78A_ | Alignment | not modelled | 97.4 | 14 | **PDB header:**structural genomics, unknown function<br>**Chain:** A: **PDB Molecule:**hypothetical protein smu.776;<br>**PDBTitle:** a putative sam-dependent methyltransferase from streptococcus mutans |
| 26 | c2esrB_ | Alignment | not modelled | 97.3 | 15 | **PDB header:**transferase<br>**Chain:** B: **PDB Molecule:**methyltransferase;<br>**PDBTitle:** conserved hypothetical protein- streptococcus pyogenes |
| 27 | c3axtA_ | Alignment | not modelled | 97.3 | 16 | **PDB header:**transferase<br>**Chain:** A: **PDB Molecule:**probable n(2),n(2)-dimethylguanosine trna methyltransferase<br>**PDBTitle:** complex structure of trna methyltransferase trm1 from aquifex aeolicus2 with s-adenosyl-l-methionine |
| 28 | d2ifta1 | Alignment | not modelled | 97.1 | 20 | **Fold:**S-adenosyl-L-methionine-dependent methyltransferases<br>**Superfamily:**S-adenosyl-L-methionine-dependent methyltransferases<br>**Family:**YhhF-like |

| | | | | | | |
|---|---|---|---|---|---|---|
| 29 | c3gdhC_ | Alignment | not modelled | 97.1 | 16 | **PDB header:**transferase<br>**Chain:** C: **PDB Molecule:**trimethylguanosine synthase homolog;<br>**PDBTitle:** methyltransferase domain of human trimethylguanosine synthase 1 (tgs1)2 bound to m7gtp and adenosyl-homocysteine (active form) |
| 30 | d1nv8a_ | Alignment | not modelled | 97.1 | 19 | **Fold:**S-adenosyl-L-methionine-dependent methyltransferases<br>**Superfamily:**S-adenosyl-L-methionine-dependent methyltransferases<br>**Family:**N5-glutamine methyltransferase, HemK |
| 31 | d2igta1 | Alignment | not modelled | 97.0 | 16 | **Fold:**S-adenosyl-L-methionine-dependent methyltransferases<br>**Superfamily:**S-adenosyl-L-methionine-dependent methyltransferases<br>**Family:**hypothetical RNA methyltransferase |
| 32 | c5hjmA_ | Alignment | not modelled | 97.0 | 14 | **PDB header:**transferase<br>**Chain:** A: **PDB Molecule:**trna (guanine(37)-n1)-methyltransferase trm5a;<br>**PDBTitle:** crystal structure of pyrococcus abyssi trm5a complexed with mta |
| 33 | d1wy7a1 | Alignment | not modelled | 96.9 | 16 | **Fold:**S-adenosyl-L-methionine-dependent methyltransferases<br>**Superfamily:**S-adenosyl-L-methionine-dependent methyltransferases<br>**Family:**Ta1320-like |
| 34 | d1uwva2 | Alignment | not modelled | 96.8 | 19 | **Fold:**S-adenosyl-L-methionine-dependent methyltransferases<br>**Superfamily:**S-adenosyl-L-methionine-dependent methyltransferases<br>**Family:**(Uracil-5-)-methyltransferase |
| 35 | d1uira_ | Alignment | not modelled | 96.7 | 28 | **Fold:**S-adenosyl-L-methionine-dependent methyltransferases<br>**Superfamily:**S-adenosyl-L-methionine-dependent methyltransferases<br>**Family:**Spermidine synthase |
| 36 | d1xj5a_ | Alignment | not modelled | 96.6 | 21 | **Fold:**S-adenosyl-L-methionine-dependent methyltransferases<br>**Superfamily:**S-adenosyl-L-methionine-dependent methyltransferases<br>**Family:**Spermidine synthase |
| 37 | c3ll7A_ | Alignment | not modelled | 96.6 | 18 | **PDB header:**transferase<br>**Chain:** A: **PDB Molecule:**putative methyltransferase;<br>**PDBTitle:** crystal structure of putative methyltransferase pg_1098 from2 porphyromonas gingivalis w83 |
| 38 | c6bq6B_ | Alignment | not modelled | 96.6 | 25 | **PDB header:**transferase<br>**Chain:** B: **PDB Molecule:**thermospermine synthase;<br>**PDBTitle:** crystal structure of medicago truncatula thermospermine synthase2 (mttsps) in complex with thermospermine |
| 39 | c2q41D_ | Alignment | not modelled | 96.5 | 22 | **PDB header:**transferase<br>**Chain:** D: **PDB Molecule:**spermidine synthase 1;<br>**PDBTitle:** ensemble refinement of the protein crystal structure of spermidine2 synthase from arabidopsis thaliana gene at1g23820 |
| 40 | c3q87B_ | Alignment | not modelled | 96.5 | 15 | **PDB header:**transferase activator/transferase<br>**Chain:** B: **PDB Molecule:**n6 adenine specific dna methylase;<br>**PDBTitle:** structure of e. cuniculi mtq2-trm112 complex responible for the2 methylation of erf1 translation termination factor |
| 41 | c6zxyB_ | Alignment | not modelled | 96.4 | 15 | **PDB header:**rna binding protein<br>**Chain:** B: **PDB Molecule:**trna (guanine(10)-n2)-dimethyltransferase;<br>**PDBTitle:** structure of archaeoglobus fulgidus trm11 m2g10 trna methyltransferase2 enzyme |
| 42 | c5xj2C_ | Alignment | not modelled | 96.3 | 15 | **PDB header:**transferase/rna<br>**Chain:** C: **PDB Molecule:**uncharacterized rna methyltransferase sp_1029;<br>**PDBTitle:** structure of sprlmcd with u747 rna |
| 43 | c6pbdA_ | Alignment | not modelled | 96.3 | 27 | **PDB header:**transferase/dna<br>**Chain:** A: **PDB Molecule:**modification methylase ccrmi;<br>**PDBTitle:** dna n6-adenine methyltransferase ccrm in complex with double-stranded2 dna oligonucleotide containing its recognition sequence gaatc |
| 44 | c1uwvA_ | Alignment | not modelled | 96.3 | 21 | **PDB header:**transferase<br>**Chain:** A: **PDB Molecule:**23s rrna (uracil-5-)-methyltransferase ruma;<br>**PDBTitle:** crystal structure of ruma, the iron-sulfur cluster2 containing e. coli 23s ribosomal rna 5-methyluridine3 methyltransferase |
| 45 | d2dula1 | Alignment | not modelled | 96.3 | 18 | **Fold:**S-adenosyl-L-methionine-dependent methyltransferases<br>**Superfamily:**S-adenosyl-L-methionine-dependent methyltransferases<br>**Family:**TRM1-like |
| 46 | c6h2uA_ | Alignment | not modelled | 96.2 | 17 | **PDB header:**transferase<br>**Chain:** A: **PDB Molecule:**methyltransferase-like protein 5;<br>**PDBTitle:** crystal structure of human mettl5-trmt112 complex, the 18s rrna2 m6a1832 methyltransferase at 1.6a resolution |
| 47 | c3bwbA_ | Alignment | not modelled | 96.1 | 27 | **PDB header:**transferase<br>**Chain:** A: **PDB Molecule:**spermidine synthase;<br>**PDBTitle:** crystal structure of the apo form of spermidine synthase from2 trypanosoma cruzi at 2.5 a resolution |
| 48 | d2pkwa1 | Alignment | not modelled | 96.0 | 19 | **Fold:**S-adenosyl-L-methionine-dependent methyltransferases<br>**Superfamily:**S-adenosyl-L-methionine-dependent methyltransferases<br>**Family:**YhiQ-like |
| 49 | d2oyra1 | Alignment | not modelled | 96.0 | 17 | **Fold:**S-adenosyl-L-methionine-dependent methyltransferases<br>**Superfamily:**S-adenosyl-L-methionine-dependent methyltransferases<br>**Family:**YhiQ-like |
| 50 | c3c6kC_ | Alignment | not modelled | 95.9 | 16 | **PDB header:**transferase<br>**Chain:** C: **PDB Molecule:**spermine synthase;<br>**PDBTitle:** crystal structure of human spermine synthase in complex2 with spermidine and 5-methylthioadenosine |
| 51 | c3tmaA_ | Alignment | not modelled | 95.9 | 16 | **PDB header:**transferase<br>**Chain:** A: **PDB Molecule:**methyltransferase;<br>**PDBTitle:** crystal structure of trmn from thermus thermophilus |
| 52 | d1iy9a_ | Alignment | not modelled | 95.8 | 19 | **Fold:**S-adenosyl-L-methionine-dependent methyltransferases<br>**Superfamily:**S-adenosyl-L-methionine-dependent methyltransferases<br>**Family:**Spermidine synthase |
| | | | | | | **Fold:**S-adenosyl-L-methionine-dependent methyltransferases |

| | | | | | | |
|---|---|---|---|---|---|---|
| 53 | d1booa_ | Alignment | not modelled | 95.8 | 24 | **Superfamily:**S-adenosyl-L-methionine-dependent methyltransferases<br>**Family:**Type II DNA methylase |
| 54 | d1g60a_ | Alignment | not modelled | 95.8 | 12 | **Fold:**S-adenosyl-L-methionine-dependent methyltransferases<br>**Superfamily:**S-adenosyl-L-methionine-dependent methyltransferases<br>**Family:**Type II DNA methylase |
| 55 | c2qm3A_ | Alignment | not modelled | 95.7 | 17 | **PDB header:**transferase<br>**Chain:** A: **PDB Molecule:**predicted methyltransferase;<br>**PDBTitle:** crystal structure of a predicted methyltransferase from pyrococcus2 furiosus |
| 56 | c3bt7A_ | Alignment | not modelled | 95.7 | 13 | **PDB header:**transferase/rna<br>**Chain:** A: **PDB Molecule:**trna (uracil-5-)-methyltransferase;<br>**PDBTitle:** structure of e. coli 5-methyluridine methyltransferase trma in complex2 with 19 nucleotide t-arm analogue |
| 57 | c3lbyA_ | Alignment | not modelled | 95.6 | 11 | **PDB header:**transferase<br>**Chain:** A: **PDB Molecule:**putative uncharacterized protein smu.1697c;<br>**PDBTitle:** crystal structure of smu.1697c, a putative methyltransferase from2 streptococcus mutans in complex with sah |
| 58 | c1wg8B_ | Alignment | not modelled | 95.4 | 27 | **PDB header:**transferase<br>**Chain:** B: **PDB Molecule:**predicted s-adenosylmethionine-dependent<br>**PDBTitle:** crystal structure of a predicted s-adenosylmethionine-2 dependent methyltransferase tt1512 from thermus3 thermophilus hb8. |
| 59 | c2zifB_ | Alignment | not modelled | 95.4 | 36 | **PDB header:**transferase<br>**Chain:** B: **PDB Molecule:**putative modification methylase;<br>**PDBTitle:** crystal structure of ttha0409, putative dna modification methylase2 from thermus thermophilus hb8- complexed with s-adenosyl-l-methionine |
| 60 | d1inla_ | Alignment | not modelled | 95.3 | 21 | **Fold:**S-adenosyl-L-methionine-dependent methyltransferases<br>**Superfamily:**S-adenosyl-L-methionine-dependent methyltransferases<br>**Family:**Spermidine synthase |
| 61 | c3grrA_ | Alignment | not modelled | 95.2 | 24 | **PDB header:**transferase<br>**Chain:** A: **PDB Molecule:**dimethyladenosine transferase;<br>**PDBTitle:** crystal structure of the complex between s-adenosyl homocysteine and2 methanocaldococcus jannaschi dim1. |
| 62 | c6qmmA_ | Alignment | not modelled | 95.2 | 19 | **PDB header:**transferase<br>**Chain:** A: **PDB Molecule:**polyamine aminopropyltransferase;<br>**PDBTitle:** crystal structure of synecochoccus spermidine synthase in complex with2 putrescine, spermidine and mta |
| 63 | d2b2ca1 | Alignment | not modelled | 95.2 | 28 | **Fold:**S-adenosyl-L-methionine-dependent methyltransferases<br>**Superfamily:**S-adenosyl-L-methionine-dependent methyltransferases<br>**Family:**Spermidine synthase |
| 64 | d1zq9a1 | Alignment | not modelled | 95.1 | 23 | **Fold:**S-adenosyl-L-methionine-dependent methyltransferases<br>**Superfamily:**S-adenosyl-L-methionine-dependent methyltransferases<br>**Family:**rRNA adenine dimethylase-like |
| 65 | c2hteB_ | Alignment | not modelled | 95.1 | 25 | **PDB header:**transferase<br>**Chain:** B: **PDB Molecule:**spermidine synthase;<br>**PDBTitle:** the crystal structure of spermidine synthase from p. falciparum in2 complex with 5'-methylthioadenosine |
| 66 | c3lpmA_ | Alignment | not modelled | 95.1 | 16 | **PDB header:**transferase<br>**Chain:** A: **PDB Molecule:**putative methyltransferase;<br>**PDBTitle:** crystal structure of putative methyltransferase small domain protein2 from listeria monocytogenes |
| 67 | c2ozvA_ | Alignment | not modelled | 95.0 | 15 | **PDB header:**transferase<br>**Chain:** A: **PDB Molecule:**hypothetical protein atu0636;<br>**PDBTitle:** crystal structure of a predicted o-methyltransferase, protein atu6362 from agrobacterium tumefaciens. |
| 68 | c3fydA_ | Alignment | not modelled | 95.0 | 24 | **PDB header:**transferase<br>**Chain:** A: **PDB Molecule:**probable dimethyladenosine transferase;<br>**PDBTitle:** crystal structure of dim1 from the thermophilic archeon,2 methanocaldococcus jannaschi |
| 69 | c2yx1A_ | Alignment | not modelled | 94.9 | 18 | **PDB header:**transferase<br>**Chain:** A: **PDB Molecule:**hypothetical protein mj0883;<br>**PDBTitle:** crystal structure of m.jannaschii trna m1g37 methyltransferase |
| 70 | c4jxjA_ | Alignment | not modelled | 94.8 | 17 | **PDB header:**transferase<br>**Chain:** A: **PDB Molecule:**ribosomal rna small subunit methyltransferase a;<br>**PDBTitle:** crystal structure of ribosomal rna small subunit methyltransferase a2 from rickettsia bellii determined by iodide sad phasing |
| 71 | c6h1dA_ | Alignment | not modelled | 94.7 | 18 | **PDB header:**gene regulation<br>**Chain:** A: **PDB Molecule:**hemk methyltransferase family member 2;<br>**PDBTitle:** crystal structure of c21orf127-trmt112 in complex with sah |
| 72 | c1nw6A_ | Alignment | not modelled | 94.6 | 18 | **PDB header:**transferase<br>**Chain:** A: **PDB Molecule:**modification methylase rsri;<br>**PDBTitle:** structure of the beta class n6-adenine dna methyltransferase rsri2 bound to sinefungin |
| 73 | c4zcfA_ | Alignment | not modelled | 94.3 | 21 | **PDB header:**hydrolase-dna complex<br>**Chain:** A: **PDB Molecule:**restriction endonuclease ecop15i, modification subunit;<br>**PDBTitle:** structural basis of asymmetric dna methylation and atp-triggered long-2 range diffusion by ecop15i |
| 74 | c2pssC_ | Alignment | not modelled | 94.1 | 25 | **PDB header:**transferase<br>**Chain:** C: **PDB Molecule:**spermidine synthase;<br>**PDBTitle:** the structure of plasmodium falciparum spermidine synthase in its apo-2 form |
| 75 | d1eg2a_ | Alignment | not modelled | 94.0 | 18 | **Fold:**S-adenosyl-L-methionine-dependent methyltransferases<br>**Superfamily:**S-adenosyl-L-methionine-dependent methyltransferases<br>**Family:**Type II DNA methylase |
| 76 | c5hfjF_ | Alignment | not modelled | 93.9 | 24 | **PDB header:**dna binding protein<br>**Chain:** F: **PDB Molecule:**adenine specific dna methyltransferase (dpna);<br>**PDBTitle:** crystal structure of m1.hpyavi-sam complex |

| | | | | | | |
|---|---|---|---|---|---|---|
| 77 | c2r6zA_ | Alignment | not modelled | 93.9 | 17 | **PDB header:**transferase<br>**Chain:** A: **PDB Molecule:**upf0341 protein in rsp 3' region;<br>**PDBTitle:** crystal structure of the sam-dependent methyltransferase ngo1261 from2 neisseria gonorrhoeae, northeast structural genomics consortium3 target ngr48 |
| 78 | c6k0wD_ | Alignment | not modelled | 93.8 | 20 | **PDB header:**dna binding protein<br>**Chain:** D: **PDB Molecule:**adenine specific dna methyltransferase (mod);<br>**PDBTitle:** dna methyltransferase in complex with sinefungin |
| 79 | c3uzuA_ | Alignment | not modelled | 93.8 | 23 | **PDB header:**transferase<br>**Chain:** A: **PDB Molecule:**ribosomal rna small subunit methyltransferase a;<br>**PDBTitle:** the structure of the ribosomal rna small subunit methyltransferase a2 from burkholderia pseudomallei |
| 80 | c6k0wB_ | Alignment | not modelled | 93.7 | 21 | **PDB header:**dna binding protein<br>**Chain:** B: **PDB Molecule:**adenine specific dna methyltransferase (mod);<br>**PDBTitle:** dna methyltransferase in complex with sinefungin |
| 81 | d1ne2a_ | Alignment | not modelled | 93.5 | 18 | **Fold:**S-adenosyl-L-methionine-dependent methyltransferases<br>**Superfamily:**S-adenosyl-L-methionine-dependent methyltransferases<br>**Family:**Ta1320-like |
| 82 | c4bluB_ | Alignment | not modelled | 93.5 | 15 | **PDB header:**transferase<br>**Chain:** B: **PDB Molecule:**ribosomal rna large subunit methyltransferase j;<br>**PDBTitle:** crystal structure of escherichia coli 23s rrna (a2030-n6)-2 methyltransferase rlmj |
| 83 | c6nvmA_ | Alignment | not modelled | 93.4 | 19 | **PDB header:**transferase<br>**Chain:** A: **PDB Molecule:**rrna adenine n-6-methyltransferase;<br>**PDBTitle:** crystal structure of 23s rrna methyltransferase erme |
| 84 | c3tqsB_ | Alignment | not modelled | 93.3 | 22 | **PDB header:**transferase<br>**Chain:** B: **PDB Molecule:**ribosomal rna small subunit methyltransferase a;<br>**PDBTitle:** structure of the dimethyladenosine transferase (ksga) from coxiella2 burnetii |
| 85 | c6ifsB_ | Alignment | not modelled | 93.3 | 22 | **PDB header:**transferase<br>**Chain:** B: **PDB Molecule:**ribosomal rna small subunit methyltransferase a;<br>**PDBTitle:** ksga from bacillus subtilis 168 |
| 86 | c6j27D_ | Alignment | not modelled | 93.3 | 15 | **PDB header:**transferase<br>**Chain:** D: **PDB Molecule:**n(4)-bis(aminopropyl)spermidine synthase;<br>**PDBTitle:** crystal structure of the branched-chain polyamine synthase from2 thermus thermophilus (tth-bpsa) in complex with n4-3 aminopropylspermidine and 5'-methylthioadenosine |
| 87 | c3fuxB_ | Alignment | not modelled | 93.2 | 19 | **PDB header:**transferase<br>**Chain:** B: **PDB Molecule:**dimethyladenosine transferase;<br>**PDBTitle:** t. thermophilus 16s rrna a1518 and a1519 methyltransferase (ksga) in2 complex with 5'-methylthioadenosine in space group p212121 |
| 88 | c3gjyA_ | Alignment | not modelled | 93.0 | 17 | **PDB header:**transferase<br>**Chain:** A: **PDB Molecule:**spermidine synthase;<br>**PDBTitle:** crystal structure of a probable spermidine synthase from2 corynebacterium glutamicum atcc 13032 |
| 89 | d2frna1 | Alignment | not modelled | 92.8 | 15 | **Fold:**S-adenosyl-L-methionine-dependent methyltransferases<br>**Superfamily:**S-adenosyl-L-methionine-dependent methyltransferases<br>**Family:**Met-10+ protein-like |
| 90 | c3fteA_ | Alignment | not modelled | 92.5 | 22 | **PDB header:**transferase/rna<br>**Chain:** A: **PDB Molecule:**dimethyladenosine transferase;<br>**PDBTitle:** crystal structure of a. aeolicus ksga in complex with rna |
| 91 | d2oo3a1 | Alignment | not modelled | 92.4 | 17 | **Fold:**S-adenosyl-L-methionine-dependent methyltransferases<br>**Superfamily:**S-adenosyl-L-methionine-dependent methyltransferases<br>**Family:**LPG1296-like |
| 92 | c3a26A_ | Alignment | not modelled | 92.2 | 15 | **PDB header:**transferase<br>**Chain:** A: **PDB Molecule:**uncharacterized protein ph0793;<br>**PDBTitle:** crystal structure of p. horikoshii tyw2 in complex with2 mesado |
| 93 | d2f8la1 | Alignment | not modelled | 92.2 | 20 | **Fold:**S-adenosyl-L-methionine-dependent methyltransferases<br>**Superfamily:**S-adenosyl-L-methionine-dependent methyltransferases<br>**Family:**N-6 DNA Methylase-like |
| 94 | c3a27A_ | Alignment | not modelled | 92.1 | 11 | **PDB header:**transferase<br>**Chain:** A: **PDB Molecule:**uncharacterized protein mj1557;<br>**PDBTitle:** crystal structure of m. jannaschii tyw2 in complex with2 adomet |
| 95 | d1qyra_ | Alignment | not modelled | 91.9 | 20 | **Fold:**S-adenosyl-L-methionine-dependent methyltransferases<br>**Superfamily:**S-adenosyl-L-methionine-dependent methyltransferases<br>**Family:**rRNA adenine dimethylase-like |
| 96 | c3tkaA_ | Alignment | not modelled | 91.8 | 26 | **PDB header:**transferase<br>**Chain:** A: **PDB Molecule:**ribosomal rna small subunit methyltransferase h;<br>**PDBTitle:** crystal structure and solution saxs of methyltransferase rsmh from2 e.coli |
| 97 | d1dusa_ | Alignment | not modelled | 91.7 | 16 | **Fold:**S-adenosyl-L-methionine-dependent methyltransferases<br>**Superfamily:**S-adenosyl-L-methionine-dependent methyltransferases<br>**Family:**Hypothetical protein MJ0882 |
| 98 | c5wcjA_ | Alignment | not modelled | 91.6 | 15 | **PDB header:**transferase<br>**Chain:** A: **PDB Molecule:**methyltransferase-like protein 13;<br>**PDBTitle:** crystal structure of human methyltransferase-like protein 13 in2 complex with sah |
| 99 | c3evzA_ | Alignment | not modelled | 91.6 | 13 | **PDB header:**transferase<br>**Chain:** A: **PDB Molecule:**methyltransferase;<br>**PDBTitle:** crystal structure of methyltransferase from pyrococcus furiosus |
| 100 | c3dmgA_ | Alignment | not modelled | 91.3 | 13 | **PDB header:**transferase<br>**Chain:** A: **PDB Molecule:**probable ribosomal rna small subunit methyltransferase;<br>**PDBTitle:** t. thermophilus 16s rrna n2 g1207 methyltransferase (rsmc) in complex2 with adohcy |

| | | | | | | |
|---|---|---|---|---|---|---|
| 101 | d1mjfa_ | Alignment | not modelled | 91.1 | 29 | **Fold:**S-adenosyl-L-methionine-dependent methyltransferases **Superfamily:**S-adenosyl-L-methionine-dependent methyltransferases **Family:**Spermidine synthase |
| 102 | d2o07a1 | Alignment | not modelled | 91.0 | 18 | **Fold:**S-adenosyl-L-methionine-dependent methyltransferases **Superfamily:**S-adenosyl-L-methionine-dependent methyltransferases **Family:**Spermidine synthase |
| 103 | c5yacA_ | Alignment | not modelled | 90.9 | 14 | **PDB header:**rna binding protein **Chain:** A: **PDB Molecule:**trna (guanine(37)-n1)-methyltransferase trm5b; **PDBTitle:** crystal structure of wt trm5b from pyrococcus abyssi |
| 104 | d2b3ta1 | Alignment | not modelled | 90.2 | 19 | **Fold:**S-adenosyl-L-methionine-dependent methyltransferases **Superfamily:**S-adenosyl-L-methionine-dependent methyltransferases **Family:**N5-glutamine methyltransferase, HemK |
| 105 | c1m6yA_ | Alignment | not modelled | 90.2 | 21 | **PDB header:**transferase **Chain:** A: **PDB Molecule:**s-adenosyl-methyltransferase mraw; **PDBTitle:** crystal structure analysis of tm0872, a putative sam-dependent2 methyltransferase, complexed with sah |
| 106 | c6b92A_ | Alignment | not modelled | 89.7 | 13 | **PDB header:**transferase **Chain:** A: **PDB Molecule:**u6 small nuclear rna (adenine-(43)-n(6))-methyltransferase; **PDBTitle:** crystal structure of the n-terminal domain of human mettl16 in complex2 with sah |
| 107 | c6qe6A_ | Alignment | not modelled | 88.2 | 5 | **PDB header:**transferase **Chain:** A: **PDB Molecule:**trna (adenine(22)-n(1))-methyltransferase; **PDBTitle:** structure of m. capricolum trmk in complex with the natural cofactor2 product s-adenosyl-homocysteine (sah) |
| 108 | c1aqjB_ | Alignment | not modelled | 88.1 | 19 | **PDB header:**methyltransferase **Chain:** B: **PDB Molecule:**adenine-n6-dna-methyltransferase taqi; **PDBTitle:** structure of adenine-n6-dna-methyltransferase taqi |
| 109 | c2vs1A_ | Alignment | not modelled | 87.8 | 18 | **PDB header:**transferase **Chain:** A: **PDB Molecule:**uncharacterized rna methyltransferase pyrab10780; **PDBTitle:** the crystal structure of pyrococcus abyssi trna (uracil-54, c5)-2 methyltransferase in complex with s-adenosyl-l-homocysteine |
| 110 | d1dcta_ | Alignment | not modelled | 87.5 | 13 | **Fold:**S-adenosyl-L-methionine-dependent methyltransferases **Superfamily:**S-adenosyl-L-methionine-dependent methyltransferases **Family:**C5 cytosine-specific DNA methylase, DCM |
| 111 | d1qama_ | Alignment | not modelled | 87.5 | 20 | **Fold:**S-adenosyl-L-methionine-dependent methyltransferases **Superfamily:**S-adenosyl-L-methionine-dependent methyltransferases **Family:**rRNA adenine dimethylase-like |
| 112 | d2h00a1 | Alignment | not modelled | 87.0 | 11 | **Fold:**S-adenosyl-L-methionine-dependent methyltransferases **Superfamily:**S-adenosyl-L-methionine-dependent methyltransferases **Family:**Methyltransferase 10 domain |
| 113 | c4dzrA_ | Alignment | not modelled | 86.6 | 18 | **PDB header:**transferase **Chain:** A: **PDB Molecule:**protein-(glutamine-n5) methyltransferase, release factor- **PDBTitle:** the crystal structure of protein-(glutamine-n5) methyltransferase2 (release factor-specific) from alicyclobacillus acidocaldarius subsp.3 acidocaldarius dsm 446 |
| 114 | d1yuba_ | Alignment | not modelled | 85.4 | 17 | **Fold:**S-adenosyl-L-methionine-dependent methyltransferases **Superfamily:**S-adenosyl-L-methionine-dependent methyltransferases **Family:**rRNA adenine dimethylase-like |
| 115 | c6q56C_ | Alignment | not modelled | 85.1 | 12 | **PDB header:**rna binding protein **Chain:** C: **PDB Molecule:**trna (adenine(22)-n(1))-methyltransferase; **PDBTitle:** crystal structure of the b. subtilis m1a22 trna methyltransferase trmk |
| 116 | d1o9ga_ | Alignment | not modelled | 84.4 | 24 | **Fold:**S-adenosyl-L-methionine-dependent methyltransferases **Superfamily:**S-adenosyl-L-methionine-dependent methyltransferases **Family:**rRNA methyltransferase AviRa |
| 117 | c2yxlA_ | Alignment | not modelled | 82.7 | 13 | **PDB header:**transferase **Chain:** A: **PDB Molecule:**450aa long hypothetical fmu protein; **PDBTitle:** crystal structure of ph0851 |
| 118 | d1m6ya2 | Alignment | not modelled | 82.3 | 19 | **Fold:**S-adenosyl-L-methionine-dependent methyltransferases **Superfamily:**S-adenosyl-L-methionine-dependent methyltransferases **Family:**MraW-like putative methyltransferases |
| 119 | c1g38A_ | Alignment | not modelled | 81.4 | 25 | **PDB header:**transferase/dna **Chain:** A: **PDB Molecule:**modification methylase taqi; **PDBTitle:** adenine-specific methyltransferase m. taq i/dna complex |
| 120 | d1sqga2 | Alignment | not modelled | 81.2 | 14 | **Fold:**S-adenosyl-L-methionine-dependent methyltransferases **Superfamily:**S-adenosyl-L-methionine-dependent methyltransferases **Family:**NOL1/NOP2/sun |
