## Supplemental for "Evolution of a Plasmid Regulatory Circuit Ameliorates Plasmid Fitness Cost"

**INDEX**

**Table**

### **Table S1. Evolved pBP136Km genotypes.**

Base pair locations of deleted regions are derived from NCBI genome for pBP136Km (accession number NZ_OR146256.1).

| **Experimental Setup** | **Deletion Size** | **Deletion Genotype** | **Deleted Region** | **Deletion Details** |
| --- | --- | --- | --- | --- |
| Clone isolated after mating of pBP136Km into nalidixic acid resistant K-12 | 168 bp | *[upf31.0]* | 16122-16289 | *upf31.0* (bp 311-478 deleted in 675 bp gene) |
| Clone Day 5A isolated from day 5 of rifampicin resistant K-12(pBP136Km) persistence assay | 1859 bp | *[trbP], upf30.5, upf31.0, [parA]* | 14676-16534 | *trbP* (bp 165-699 deleted in 699 bp gene)  *upf30.5* & *upf31.0* entirely deleted.  *parA* (bp 806-852 deleted in 852 bp gene) |
| Clone Day 5B isolated from day 5 of rifampicin resistant K-12(pBP136Km) persistence assay | 871 bp | *[upf30.5]-[upf31.0]* | 15514-16385 | *Upf30.5* (bp 289-432 deleted in 432 bp gene)  *Upf31.0* (bp 1-572 deleted in 675 bp gene) |
| Clone Day 5C isolated from day 5 of rifampicin resistant K-12(pBP136Km) persistence assay | Same as above | Same as above | Same as above | Same as above |
| Clone A post-electroporation of pBP136Km into *E. coli* BW25113 | 2 bp | Intergenic (promoter region) | 15757-15758 | *upf31* promoter |
| Clone B post-electroporation of pBP136Km into *E. coli* BW25113 | 91 bp | *[upf31.0]* | 16042-16131 | *Upf31.0* (bp 231-320 deleted in 675 bp gene) |

### **Table S2. Upf31 amino acid differences between pBP136Km and pAKD1 or pALTS29.** Non-conservative substitutions in bold. Differences to Upf31/pBP136Km that are conserved in pAKD1 and pALTS29 are italicized.

| **pBP136Km -> pAKD1** | **pBP136Km -> pALTS29** |
| --- | --- |
| *I47V* | *I47V* |
| *I51V* | *I51V* |
| R60Q |  |
| *Y70H* | *Y70H* |
| ***D76A*** | ***D76A*** |
|  | L78R |
|  | Q82P |
|  | S86T |
| ***A91V*** | ***A91V*** |
| *Y108H* | *Y108H* |
| ***R124G*** | ***R124G*** |
| *K174E* | *K174E* |
| *I202L* | *I202L* |

### **Table S3. *xylE* reporter gene activity from upf31p and effect of Upf31 on expression.**

| **Plasmid(s)** | **rate dAB/min** | **[protein]** | **XylE units** | **Mean** | **SD** | **Two-sample independent T-test** |
| --- | --- | --- | --- | --- | --- | --- |
| pGCMT1 (no promoter) | 0 | 0.025 | 0 | 0.088 | 0.076 |  |
| pGCMT1 (no promoter) | 0.0057 | 0.043 | 0.133 |  |  |  |
| pGCMT1 (no promoter) | 0.0061 | 0.047 | 0.130 |  |  |  |
| pGCMT*upf3*1p | 0.52 | 0.058 | 8.966 | 7.439 | 1.660 | 0.016 |
| pGCMT*upf31*p | 0.43 | 0.056 | 7.679 |  |  |  |
| pGCMT*upf31*p | 0.38 | 0.067 | 5.672 |  |  |  |
| pGCMT*upf31*p + pBR322 | 0.24 | 0.052 | 4.616 | 4.755 | 0.121 |  |
| pGCMT*upf31*p + pBR322 | 0.26 | 0.054 | 4.815 |  |  |  |
| pGCMT*upf31*p + pBR322 | 0.29 | 0.06 | 4.833 |  |  |  |
| pGCMTupf31p + pBR*upf31* | 0.0062 | 0.049 | 0.127 | 0.094 | 0.032 | 8.996E-05 |
| pGCMTupf31p + pBR*upf31* | 0.0049 | 0.052 | 0.094 |  |  |  |
| pGCMTupf31p + pBR*upf31* | 0.0041 | 0.066 | 0.062 |  |  |  |

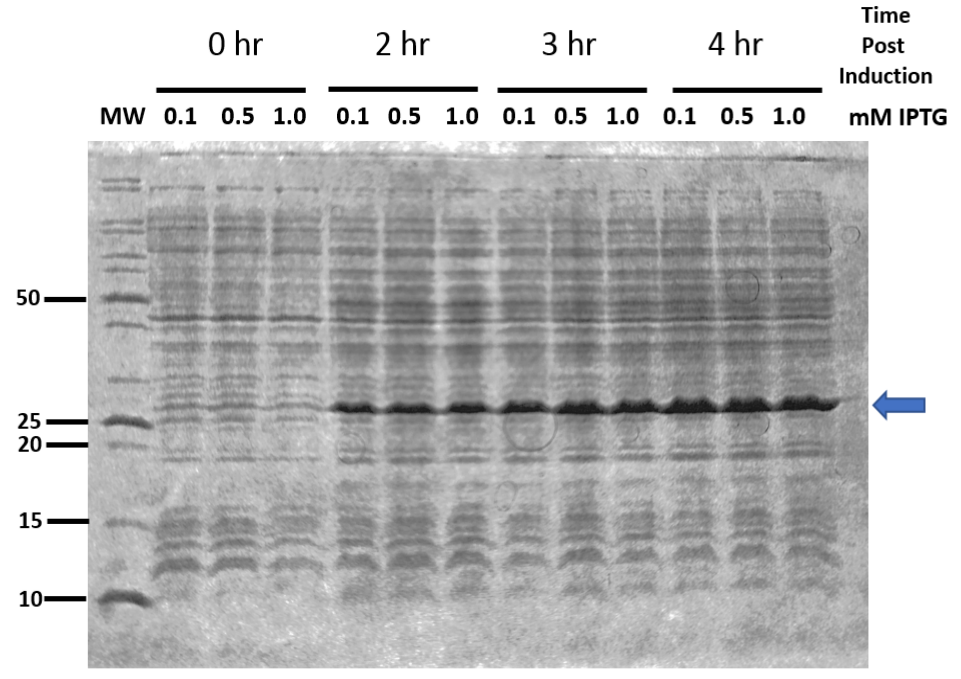

### **Figure S1. Upf31 SDS-Page protein gel**

Coomassie stained 12% SDS-PAGE demonstrating expression of upf31. MW is the protein mass standards (pertinent masses indicated left of the gel). The blue arrow indicates the position of the upf31 protein (expected mass 25.4 kDa).

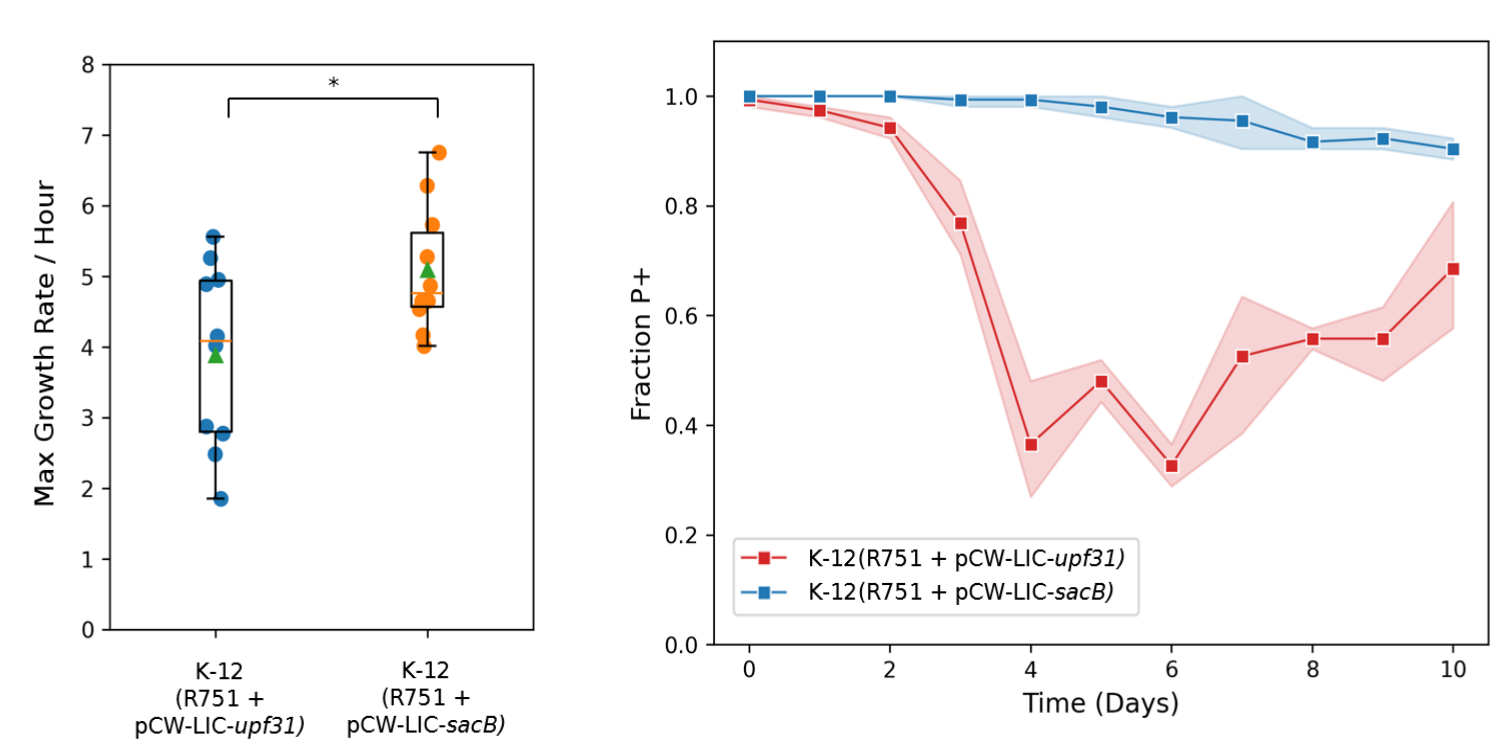

### **Figure S2. Maximum growth rates and persistence of archetype IncP plasmid R751 in *E. coli* K-12 (R751) (pCW-LIC-*upf31*) and control *E. coli* K-12 (R751) (pCW-LIC-*sacB*).**

Left: Two sample independent T-test p-value = 0.02676, *n*=10. Box is interquartile range, green triangle is mean, orange line is median, *n.s.*=p > 0.05, *=p ≤ 0.05.

Right: Lighter shades indicate 95% confidence interval. The increase in persistence after day four in the K-12(R751 + pCW-LIC-*upf31)* populations is likely due to clonal interference between evolved pCW-LIC-*upf31*. A clone isolated from day 0 showed no mutations in pCW-LIC-*upf31* or R751. Two clones (CFU A and B) from day 10 were randomly selected and sequenced via Illumina short reads. The sequencing data from both clones showed plasmid heterogenicity of pCW-LIC-*upf31* and no changes in R751*.*We cannot ascertain what proportion of cells contained which pCW-LIC-*upf31*variants, but the variants we identified are as follows: (i) CFU A contained a pCW-LIC-*upf31* variant with complete deletion of the two *tac* promoters upstream of *upf31;* (ii) CFU B contained:

- A pCW-LIC-*upf31* variant with a deletion of the last 367 base pairs of *upf31* and part of the F1 origin of replication (a secondary *oriV* on PCW-LIC).
- A pCW-LIC-*upf31* variant with a 293 base pair deletion entirely within the *upf31* coding sequence.
- A pCW-LIC-*upf31* variant with complete loss of the *lacL* gene and the entire promoter region just upstream of *upf31*.

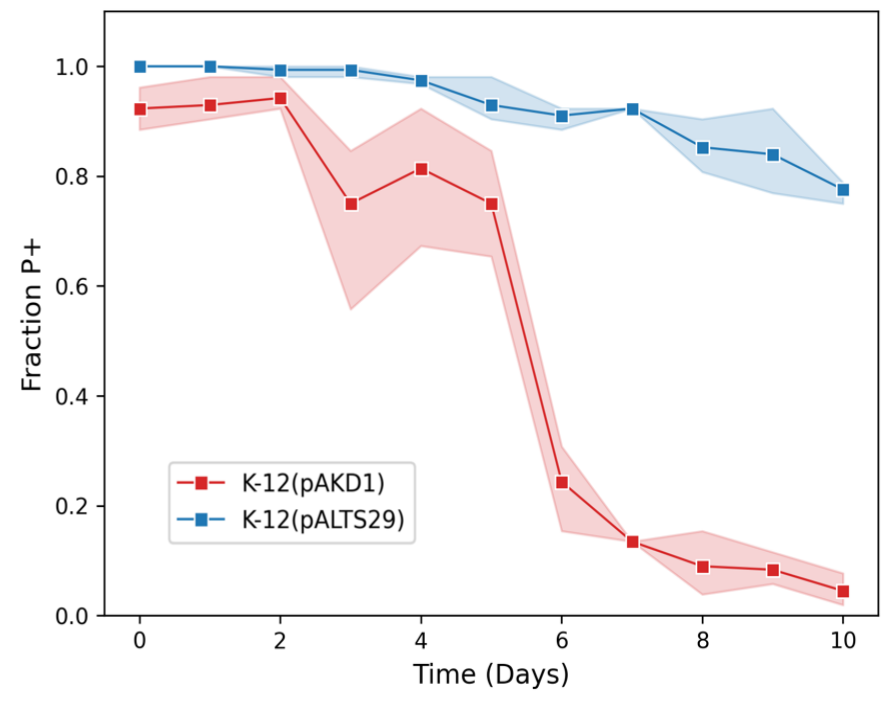

### **Figure S3. Persistence of IncP plasmids pAKD1 and pALTS29 in *E. coli* K-12.** Lighter shades indicate 95% confidence interval.

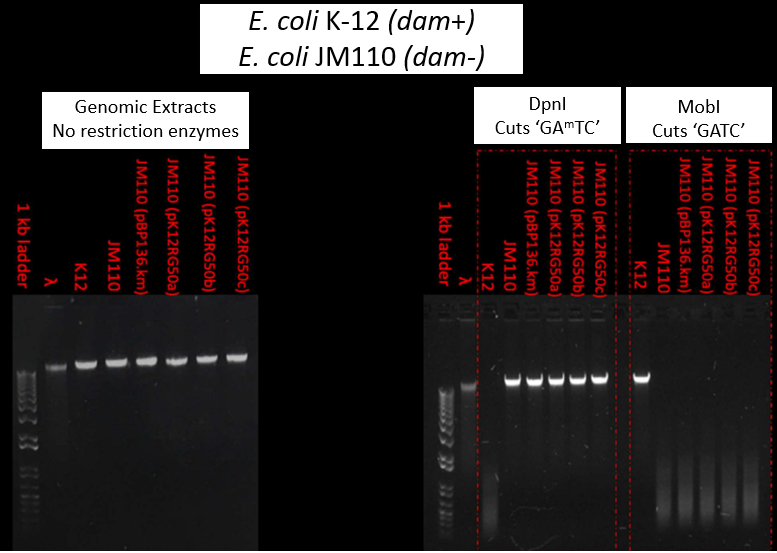

### **Figure S4. Restriction fragment length profiles of *dam*+ and *dam*- *E. coli* strains suggesting *upf31* does not encode a functional Dam methylase**

To test if *upf31* is a functional homologue to *dam*, we performed a comparative restriction enzyme assay. The Dam protein methylates the adenine base in ‘GATC’ sequences (*i.e. ‘*GA^m^TC’). We utilized two restriction enzymes, *DpnI* and *MobI*, each of which cuts GATC sequences but with different methylation specificity. *DpnI* only cuts GA^m^TC, while *MobI* cuts unmethylated GATC base pairs.

*Escherichia coli* K-12 natively encodes *dam* and has methylated GA^m^TC base pairs throughout its genome. Conversely, *E. coli* JM110 contains a *dam* knockout and contains unmethylated GATC. Thus K-12 has a *dam+* and JM110 has a *dam*- methylation profile.

The agarose gels show genomic DNA extracted from the respective strains, which appear as solid bright lines if uncut (left hand side). On the right hand side, when using enzyme *DpnI,* as expected, the DNA of K-12with its *dam+* profile is clearly cut, represented by a smear, whereas the DNA of *E. coli* JM110 is not due to its *dam-* profile; JM110 with pBP136Km and several evolved plasmid variants likewise remain uncut with *DpnI*, suggesting that *upf31* does not restore *dam+* methylation patterns.

A similar experiment was repeated, but with *MobI* instead, which cuts any unmethylated GATC sequences (far right side of the gel). Note the DNA of JM110 with its *dam-* phenotype is cut, along with that of JM110 with pBP136Km and several evolved plasmid variants. This confirms that pBP136Km with *upf31* does not restore *dam+* methylation patterns to JM110. Conversely, as the methylated GA^m^TC sequences in K-12 are impervious to *MobI* cuts the genomic DNA remained uncut.

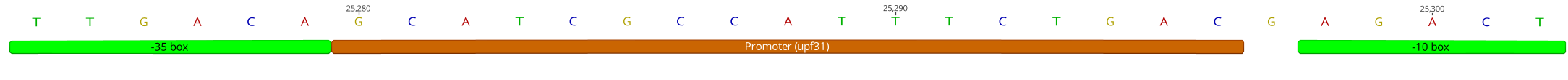

5’- TTGACAGCATCGCCATTTCTGACGAGACT-3’

### **Figure S5. Putative *upf31* promoter in pBP136Km.**

**-35	>>>>>	IVR	<<<<<**

**upf31>>**CTAGATTTAGCCGCTAAAGGCCCGGCCGGCTTCCC**TTGACA**GCATCGCCATTTCTGAC**GATGCT**GGCCAAGTCCCGATTTACTCCAGGAATTCGTTGGCGGAAACAAACCTGACAACATGAACTATGAAGAGGTGACGTC 140

**xylE>>**ATGAACAAAGGTGTAATGCGACCGGGCCATGTGCAGCTGCGTGTACTGGACATGAGCAAGGCCCTGGAACACTACGTCGAGTTGCTGGGCCTGATCGAGATGGACCGTGACGACCAGGGCCGTGTCTATCTGAAGGCTTGGACCGAAGTGGATAAGTTTTCCCTGGTGCTACGCGAGGCTGACGAGCCGGGCATGGATTTTATGGGTTTCAAGGTTGTGGATGAGGATGCTCTCCGGCAACTGGAGCGGGATCTGATGGCATATGGCTGTGCCGTTGAGCAGCTACCCGCAGGTGAACTGAACAGTTGTGGCCGGCGCGTGCGCTTCCAGGCCCCCTCCGGGCATCACTTCGAGTTGTATGCAGACAAGGAATATACTGGAAAGTGGGGTTTGAATGACGTCAATCCCGAGGCATGGCCGCGCGATCTGAAAGGTATGGCGGCTGTGCGTTTCGACCACGCCCTCATGTATGGCGACGAATTGCCGGCGACCTATGACCTGTTCACCAAGGTGCTCGGTTTCTATCTGGCCGAACAGGTGCTGGACGAAAATGGCACGCGCGTCGCCCAGTTTCTCAGTCTGTCGACCAAGGCCCACGACGTGGCCTTCATTCACCATCCGGAAAAAGGCCGCCTCCATCATGTGTCCTTCCACCTCGAAACCTGGGAAGACTTGCTTCGCGCCGCCGACCTGATCTCCATGACCGACACATCTATCGATATCGGCCCAACCCGCCACGGCCTCACTCACGGCAAGACCATCTACTTCTTCGACCCGTCCGGTAACCGCAACGAAGTGTTCTGCGGGGGAGATTACAACTACCCGGACCACAAACCGGTGACCTGGACCACCGACCAGCTGGGCAAGGCGATCTTTTACCACGACCGCATTCTCAACGAACGATTCATGACCGTGCTGACCTGATGGTCCGGTACGACTTATTGCAGAGAAAAAGCCCGGCGTTGCCGGGCTTGTTTTTCAGCGTAATGCTCTGCCAGTGTTACAACCAATTAACCAATTCTGA 1164

**aphA<<**TTAGAAAAACTCATCCAGCATCAAATGAAACTGCAATTTATTCATATCAGGATTATCAATACCATATTTTTGAAAAAGCCGTTTCTGTAATGAAGGAGAAAACTCACCGAGGCAGTTCCATAGGATGGCAAGATCCTGGTATCGGTCTGCGATTCCGACTCGTCCAACATCAATACAACCTATTAATTTCCCCTCGTCAAAAATAAGGTTATCAAGTGAGAAATCACCATGAGTGACGACTGAATCCGGTGAGAATGGCAAAAGCTTATGCATTTCTTTCCAGACTTGTTCAACAGGCCAGCCATTACGCTCGTCATCAAAATCACTCGCATCAACCAAACCGTTATTCATTCGTGATTGCGCCTGAGCGAGACGAAATACGCGATCGCTGTTAAAAGGACAATTACAAACAGGAATCGAATGCAACCGGCGCAGGAACACTGCCAGCGCATCAACAATATTTTCACCTGAATCAGGATATTCTTCTAATACCTGGAATGCTGTTTTCCCGGGGATCGCAGTGGTGAGTAACCATGCATCATCAGGAGTACGGATAAAATGCTTGATGGTCGGAAGAGGCATAAATTCCGTCAGCCAGTTTAGTCTGACCATCTCATCTGTAACATCATTGGCAACGCTACCTTTGCCATGTTTCAGAAACAACTCTGGCGCATCGGGCTTCCCATACAATCGATAGATTGTCGCACCTGATTGCCCGACATTATCGCGAGCCCATTTATACCCATATAAATCAGCATCCATGTTGGAATTTAATCGCGGCCTCGAGCAAGACGTTTCCCGTTGAATATGGCTCATAACACCCCTTGTATTACTGTTTATGTAAGCAGACAGTTTTATTGTTCATGATGATATATTTTTATCTTGTGCAATGTAACATCAGAGATTTTGAGACACAACGTGGCTTTGTTG 2094

**pSC101>>**GCCACTTTCAGTATGGATTTGGGTGTTACGGGGGGGGGTTGGTGCTTAAAATCGCGCTGAGGATTTCTGATGGTGTTAAGCGGGCGGTTTTGAGATGTAAACTCGCCCGTTTAACATAATGGATCTTGCGCGCACCGCCCGAACACCACTCGCCACAAAAAACCGCCGGAACGTCCAAAAGTACGGGTTTTGCTGCCCGCAAACGGGCTGTTCTGGTGTTGCTAGTTTGTTATCAGAATCGCAGATCCGGCTTCAGCCGGTTTGCCGGCTGAAAGCGCTATTTCTTCCAGAATTGCCATGATTTTTTCCCCACGGGAGGCGTCACTGGCTCCCGTGTTGTCGGCAGCTTTGATTCGATAAGCAGCATCGCCTGTTTCAGGCTGTCTATGTGTGACTGTTGAGCTGTAACAAGTTGTCTCAGGTGTTCAATTTCATGTTCTAGTTGCTTTGTTTTACTGGTTTCACCTGTTCTATTAGGTGTTACATGCTGTTCATCTGTTACATTGTCGATCTGTTCATGGTGAACAGCTTTGAATGCACCAAAAACTCGTAAAAGCTCTGATGTATCTATCTTTTTTACACCGTTTTCATCTGTGCATATGGACAGTTTTCCCTTTGATATGTAACGGTGAACAGTTGTTCTACTTTTGTTTGTTAGTCTTGATGCTTCACTGATAGATACAAGAGCCATAAGAACCTCAGATCCTTCCGTATTTAGCCAGTATGTTCTCTAGTGTGGTTCGTTGTTTTTGCGTGAGCCATGAGAACGAACCATTGAGATCATACTTACTTTGCATGTCACTCAAAAATTTTGCCTCAAAACTGGTGAGCTGAATTTTTGCAGTTAAAGCATCGTGTAGTGTTTTTCTTAGTCCGTTATGTAGGTAGGAATCTGATGTAATGGTTGTTGGTATTTTGTCACCATTCATTTTTATCTGGTTGTTCTCAAGTTCGGTTACGAGATCCATTTGTCTATCTAGTTCAACTTGGAAAATCAACGTATCAGTCGGGCGGCCTCGCTTATCAACCACCAATTTCATATTGCTGTAAGTGTTTAAATCTTTACTTATTGGTTTCAAAACCCATTGGTTAAGCCTTTTAAACTCATGGTAGTTATTTTCAAGCATTAACATGAACTTAAATTCATCAAGGCTAATCTCTATATTTGCCTTGTGAGTTTTCTTTTGTGTTAGTTCTTTTAATAACCACTCATAAATCCTCATAGAGTATTTGTTTTCAAAAGACTTAACATGTTCCAGATTATATTTTATGAATTTTTTTAACTGGAAAAGATAAGGCAATATCTCTTCACTAAAAACTAATTCTAATTTTTCGCTTGAGAACTTGGCATAGTTTGTCCACTGGAAAATCTCAAAGCCTTTAACCAAAGGATTCCTGATTTCCACAGTTCTCGTCATCAGCTCTCTGGTTGCTTTAGCTAATACACCATAAGCATTTTCCCTACTGATGTTCATCATCTGAGCGTATTGGTTATAAGTGAACGATACCGTCCGTTCTTTCCTTGTAGGGTTTTCAATCGTGGGGTTGAGTAGTGCCACACAGCATAAAATTAGCTTGGTTTCATGCTCCGTTAAGTCATAGCGACTAATCGCTAGTTCATTTGCTTTGAAAACAACTAATTCAGACATACATCTCAATTGGTCTAGGTGATTTTAATCACTATACCAATTGAGATGGGCTAGTCAATGATAATTACTAGTCCTTTTCCTTTGAGTTGTGGGTATCTGTAAATTCTGCTAGACCTTTGCTGGAAAACTTGTAAATTCTGCTAGACCCTCTGTAAATTCCGCTAGACCTTTGTGTGTTTTTTTTGTTTATATTCAAGTGGTTATAATTTATAGAATAAAGAAAGAATAAAAAAAGATAAAAAGAATAGATCCCAGCCCTGTGTATAACTCACTACTTTAGTCAGTTCCGCAGTATTACAAAAGGATGTCGCAAACGCTGTTTGCTCCTCTACAAAACAGACCTTAAAACCCTAAAGGCTTAAGTAGCACCCTCGCAAGCTCGGGCAAATCGCTGAATATTCCTTTTGTCTCCGACCATCAGGCACCTGAGTCGCTGTCTTTTTCGTGACATTCAGTTCGCTGCGCTCACGGCTCTGGCAGTGAATGGGGGTAAATGGCACTACAGGCGCCTTTTATGGATTCATGCAAGGAAACTACCCATAATACAAGAAAAGCCCGTCACGGGCTTCTCAGGGCGTTTTATGGCGGGTCTGCGGAGTGGTGAATCCGTTAGCGAGGTGCCGCCGGCTTCCATTCAGGTCGAGGTGGCCCGGCTCCATGCACCGCGACGCAACGCGGGGAGGCAGACAAGGTATAGGGCGGCGCCTACAATCCATGCCAACCCGTTCCATGTGCTCGCCGAGGCGGCATAAATCGCCGTGACGATCAGCGGTCCAATGATCGAAGTTAGGCTGGTAAGAGCCGCGAGCGATCCTTGAAGCTGTCCCTGATGGTCGTCATCTACCTGCCTGGACAGCATGGCCTGCAACGCGGGCATCCCGATGCCGCCGGAAGCGAGAAGAATCATAATGGGGAAGGCCATCCAGCCTCGCGTCGCGAACGCCAGCAAGACGTAGCCCAGCGCGTCGGCCGCCATGCCGGCGATAATGGCCTGCTTCTCGCCGAAACGTTTGGTGGCGGGACCAGTGACGAAGGCTTGAGCGAGGGCGTGCAAGATTCCGAATACCGCAAGCGACAGGCCGATCATCGTCGCGCTCCAGCGAAAGCGGTCCTCGCCGAAAATGACCCAGAGCGCTGCCGGCACCCTGTCCTACGAGTTGCATGATAAAGAAGACAGTCATAAGTGCGGCGACGATAGTCATGCCCCGCGCCCACCGGAAGGAGCTGACTGGGTTGAAGGCTCTCAAGGGCATCGT 4999

### **Figure S6. DNA sequence of pGCMT-upf31p.** Key regions include: *upf31* promoter (cyan), *xylE* reporter gene (yellow), *aphA* conferring kanamycin resistance (red), and the segment from pSC101 including its low copy number replicon. Directionality of *xylE* and *aphA* are shown with >> and <<. In the cyan promoter region the putative -35 and -10 regions are shown in bold and underlined. The inverted repeats of the putative Upf31 binding site are also indicated.

### **Data S1. Plasmids containing *upf31***

See ‘supplemental_data_S1.xlsx’ file

### **Data S2. PHYRE predictions of *upf31* function**

See ‘supplemental_data_S2.pdf’ file

### **Data S3. RNA-Seq results of K-12(pBP136Km) and K-12(pBP136KmΔ*upf31)***

See ‘supplemental_data_S3.xlsx’ file

### **Data S4. DAVID predictions of differentially expressed cellular pathways**

See ‘supplemental_data_S4.xlsx’ file

### **Data S5. Interactions between AlpaFold2 predicted Upf31 and pBP136-encoded proteins quantified by self-assessment ranking score**

See ‘supplemental_data_S5.xlsx’ file

### **Data S6. Upf31/R751 aligned to Upf31/pBP136Km**

See ‘supplemental_data_S6.fasta’ file
